# Systematic measurement of combination drug landscapes to predict *in vivo* treatment outcomes for tuberculosis

**DOI:** 10.1101/2021.02.03.429579

**Authors:** Jonah Larkins-Ford, Talia Greenstein, Nhi Van, Yonatan N. Degefu, Michaela C. Olson, Artem Sokolov, Bree B. Aldridge

**Affiliations:** Department of Molecular Biology and Microbiology, Tufts University School of Medicine, Boston, MA; Stuart B. Levy Center for Integrated Management of Antimicrobial Resistance, Boston, MA; Graduate School of Biomedical Sciences, Tufts University School of Medicine, Boston, MA; Laboratory of Systems Pharmacology, Harvard Medical School, Boston, MA; Department of Biomedical Engineering, Tufts University School of Engineering, Medford, MA

## Abstract

A lengthy multidrug chemotherapy is required to achieve a durable cure in tuberculosis. Variation in *Mycobacterium tuberculosis* drug response is created by the differing microenvironments in lesions, which generate different bacterial drug susceptibilities. To better realize the potential of combination therapy to shorten treatment duration, multidrug therapy design should deliberately explore the vast combination space. We face a significant scaling challenge in making systematic drug combination measurements because it is not practical to use animal models for comprehensive drug combination studies, nor are there well-validated high-throughput *in vitro* models that predict animal outcomes. We hypothesized that we could both prioritize combination therapies and quantify the predictive power of various *in vitro* models for drug development using a dataset of drug combination dose responses measured in multiple *in vitro* models. We systematically measured *M. tuberculosis* response to all 2- and 3-drug combinations among ten antibiotics in eight conditions that reproduce lesion microenvironments. Applying machine learning to this comprehensive dataset, we developed classifiers predictive of multidrug treatment outcome in a mouse model of disease relapse. We trained classifiers on multiple mouse models and identified ensembles of *in vitro* models that best describe *in vivo* treatment outcomes. Furthermore, we found that combination synergies are less important for predicting outcome than metrics of potency. Here, we map a path forward to rationally prioritize combinations for animal and clinical studies using systematic drug combination measurements with validated *in vitro* models. Our pipeline is generalizable to other difficult-to-treat diseases requiring combination therapies.

**One Sentence Summary:** Signatures of *in vitro* potency and drug interaction measurements predict combination therapy outcomes in mouse models of tuberculosis.

## Introduction

Tuberculosis (TB), caused by infection with *Mycobacterium tuberculosis* (Mtb), remains a major global health issue. In 2019, an estimated ten million people fell ill with TB, and about 1.4 million people died *(1)*. Development of shorter treatment regimens is a key part of the third pillar of the WHO End TB Strategy *(2)*. Multidrug treatment regimens were developed to treat active TB infections by shortening treatment duration, reducing disease relapse, and decreasing antibiotic resistance development *(3)*. The standard TB treatment is six to nine months of multidrug treatment with an estimated 85% cure rate *(1, 4, 5)*. The first two months of treatment (intensive, bactericidal phase) consist of four drugs (isoniazid, rifampicin, pyrazinamide, and ethambutol) that reduce sputum Mtb levels but are less effective against non-replicative bacilli *(3, 4, 6)*. The following four to seven months of treatment (continuation phase) consist of two drugs (isoniazid and rifampicin) aimed at reducing disease relapse by treating persisting bacteria that survived the intensive phase *(3, 4, 6)*. New regimens that can more efficiently treat Mtb are needed to shorten the intensive phase of treatment and reduce or eliminate the bacteria that persist and require continuation phase treatment *(4)*.

Due, in large part, to the heterogeneity of TB lesions and treatment response among the Mtb population, combination therapy is required to treat active TB. Therapies should therefore be designed as combinations of antibiotics rather than single antibiotics alone. There are many drug options for new treatment regimens using existing drugs and drugs in development *(7)*, which creates an enormous number of possible drug combinations *(5)*. Preliminary results from a Phase 3 clinical trial (“Study 31”) demonstrated that treatment could be shortened using a novel combination of existing TB antibiotics *(8, 9)*. Relatively new drugs that can target non-replicative bacteria (bedaquiline, pretomanid, delamanid, SQ109) *(10–12)* in combination with established drugs are in new, treatment-shorting regimens for multidrug resistant TB (MDR-TB) *(5, 13, 14)*.

Treatment shortening potential in Phase 2b trials *(15, 16)*) led to the Phase 3 STAND clinical trial testing the use of pretomanid with pyrazinamide and moxifloxacin, PaMZ (Table 1) *(17)*. Adding bedaquiline to PaMZ (BPaMZ, Table 1) in a Phase 2b trial *(18)* shortened culture conversion time of MDR-TB so dramatically that the STAND trial was put on permanent hold to start the Phase 3 SimpliciTB trial to evaluate BPaMZ for treating both drug-sensitive TB and MDR-TB *(9, 19)*. Together, these studies and the history of TB drug regimen design has demonstrated that there is treatment-shortening potential in the drug combination space. A critical step for developing new treatment regimens is prioritizing the thousands of drug combinations before clinical testing. However, it is not practical to evaluate thousands of combinations using the current preclinical regimen design pipeline, which combines *in vitro* and small animal studies. An efficient methodology is needed to systematically assess drug combinations and prioritize the thousands of multidrug combinations for their treatment-shorting potential.

**Table 1.**
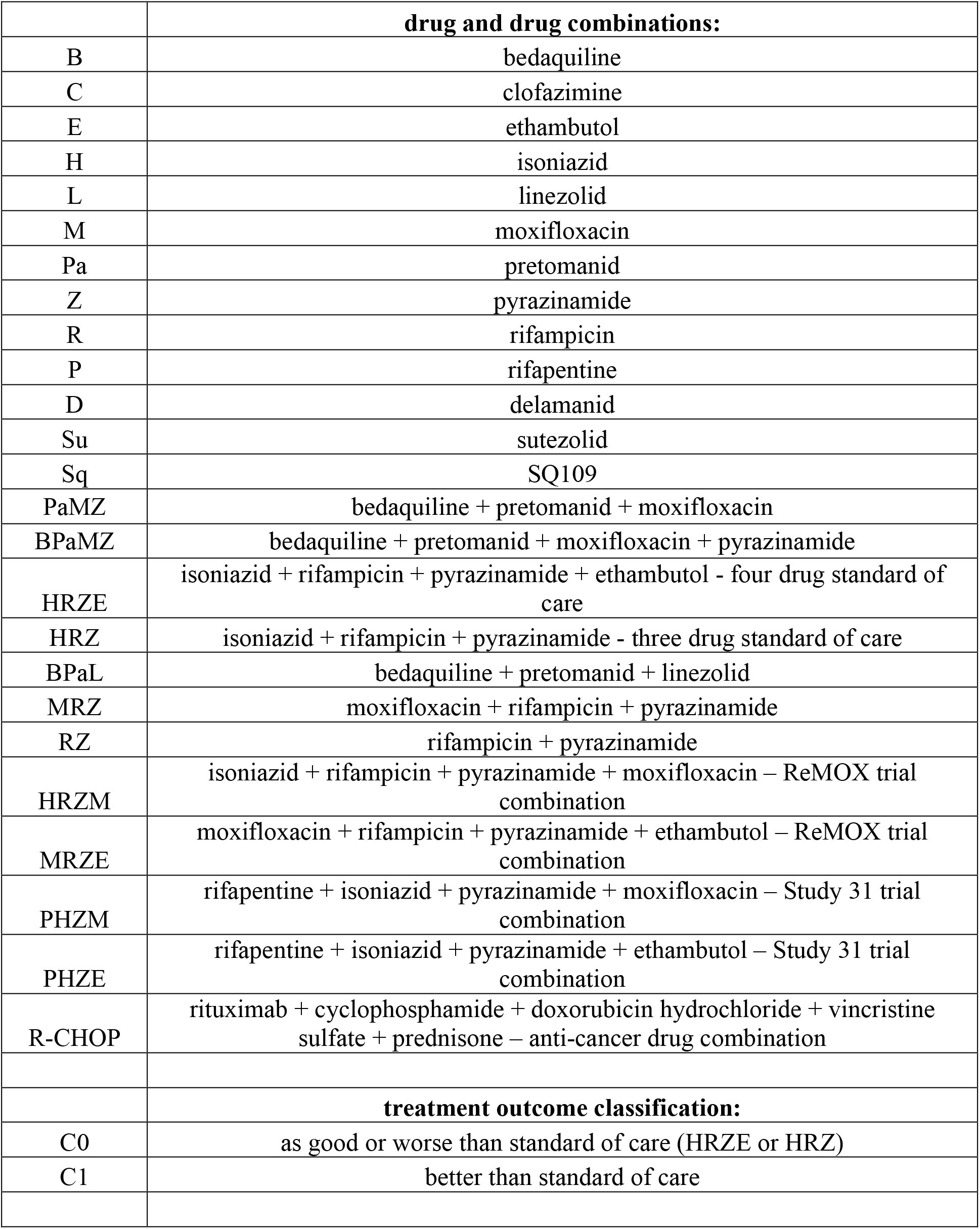
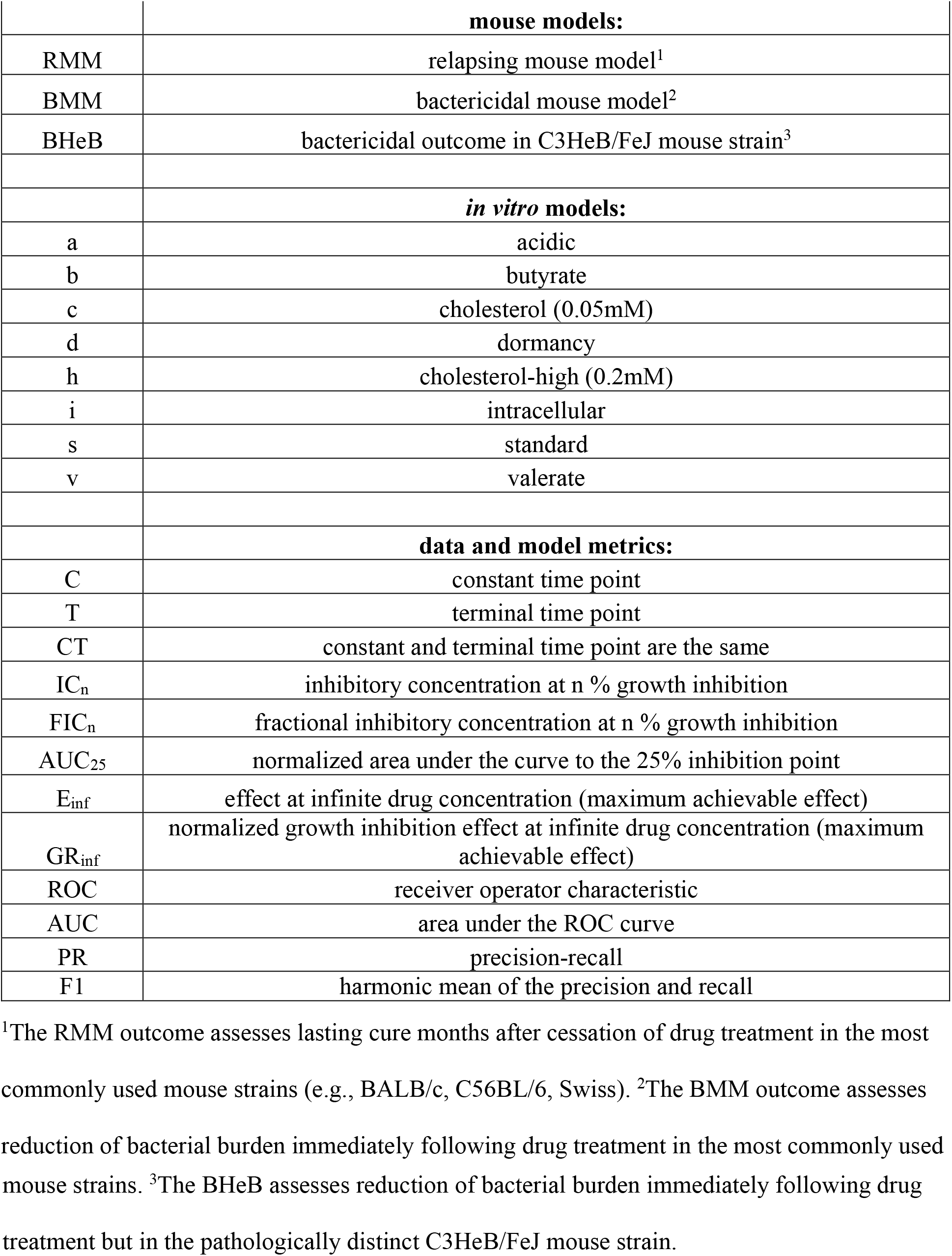
Abbreviations used in this study. Abbreviations along with brief descriptions are listed. ^1^The RMM outcome assesses lasting cure months after cessation of drug treatment in the most commonly used mouse strains (e.g., BALB/c, C56BL/6, Swiss). ^2^The BMM outcome assesses reduction of bacterial burden immediately following drug treatment in the most commonly used mouse strains. ^3^The BHeB assesses reduction of bacterial burden immediately following drug treatment but in the pathologically distinct C3HeB/FeJ mouse strain.

Animal models are critical to regimen development, and mouse models are a primary tool in multidrug therapy design *(20–24)*. Mouse strains where Mtb is primarily intracellular (e.g., BALB/c and C57BL/6) are the most widely used *(24)*. Mouse strains that form mixed lesion types (e.g., C3HeB/FeJ) are used to study drug response because the disease pathology is more humanlike, include granulomas with caseous necrotic cores *(21, 25, 26)*. Mtb drug response differs between these two types of mouse models, and both are important preclinical tools because the model-specific drug response is thought to result from the different lesion microenvironments present in each animal model *(24, 27–29)*. Despite their utility for regimen development, comprehensive drug measurements in mice are not feasible. It is only practical to perform systematic drug combination studies *in vitro*, but *in vitro* studies do not clearly map to *in vivo* outcomes *(24, 30)*. Many *in vitro* models mimic aspects of the host microenvironment encountered in the different TB lesion types. Many of these *in vitro* models are well suited for systematic drug combination studies, but none have been validated to prioritize drug combinations against preclinical animal models.

We propose to realize the potential of drug combinations to improve treatment by developing a pipeline to map *in vitro* measurement of drug response to outcomes in mouse models. Here, we utilized the efficiency of an experimental design and analysis method called DiaMOND (diagonal measurement of n-way drug interactions) *(31)* to create a compendium of drug combination responses in Mtb using multiple *in vitro* models that were designed to reproduce aspects of the environments encountered in different lesion types. Applying machine learning to this comprehensive *in vitro* dataset, we identified signatures of drug potency and interaction that could predict whether combinations would outperform the standard of care. Classifiers based on these signatures also enabled us to establish a mapping between *in vitro* models and the different mouse models, which differ in lesion type (microenvironment) and outcome. Overall, our study establishes a logistical path to optimize combination therapies for TB using systematic measurement in validated *in vitro* growth models and computational modeling.

## Results

### Drug combination compendium construction

We developed a pipeline to efficiently prioritize drug combinations early in regimen development based on drug combination measurements from *in vitro* models. Using the DiaMOND methodology *(31)*, we designed a compendium of drug combination measurements to survey informative drug-dose combinations (DiaMOND compendium). To compare *in vitro* data to treatment responses in animal models, our DiaMOND compendium focused on (A) first- and second-line agents, for which there are abundant animal data and (B) measurements in *in vitro* growth conditions that model environments encountered during infection.

Mtb encounters a diversity of environmental niches during infection that influence response to drug treatment. We aimed to model drug response by aggregating measurements from a suite of *in vitro* models. We focused on modeling factors previously shown to influence Mtb growth and/or drug response, such as different carbon sources and abundance, low pH, low oxygen tension, and the intracellular environment *(21, 30, 32–40)*. We developed or adapted eight *in vitro* models that were reproducible and scalable for systematic, high-throughput drug combination studies for this study. We varied carbon sources, with an emphasis on cholesterol and fatty acids, to model the lipid-rich environment in TB granulomas, using butyrate, valerate, cholesterol, and higher levels of cholesterol (cholesterol-high) as sole carbon sources. We used 7H9-based medium to compare against the most commonly utilized *in vitro* growth model with glycerol as a carbon source (standard). We also included *in vitro* models that mimic important factors encountered during infection: low pH (acidic), infection of J774 macrophages (intracellular), and developed a low-oxygen multi-stress model that induces dormancy using butyrate as a carbon source, sodium nitrate to respire *(41–43)*, and plate seals to limit oxygen (dormancy). The doubling times varied considerably among the models, ranging from 16h to one week (Fig. 1A). We scaled the timing of the experiments relative to the doubling time of each model so that drug response measurements would not be biased by changes in growth rate (Table S1).

**Fig. 1.**
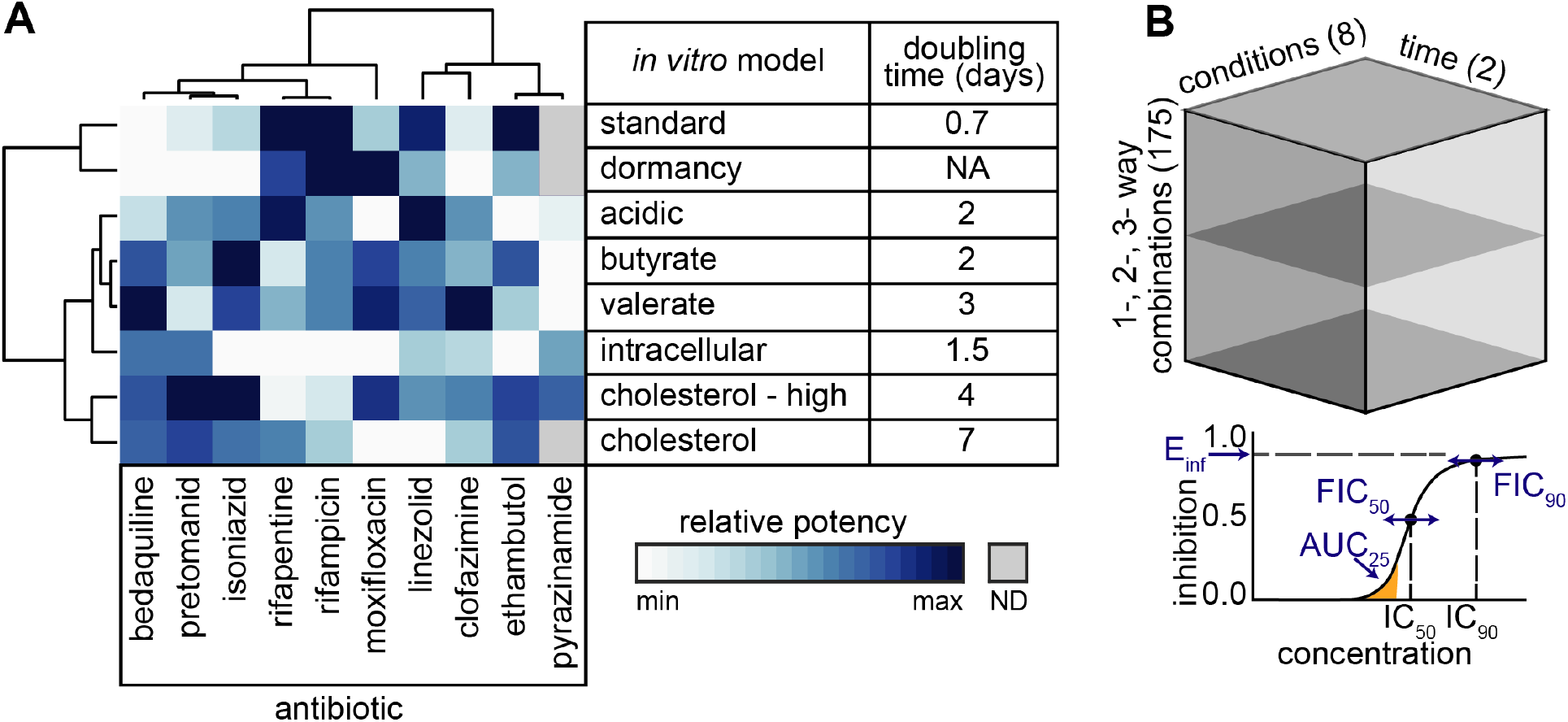
10-drug DiaMOND compendium of Mtb response to drug combination treatment. (A) Relative potencies of the ten compendium drugs in eight *in vitro* conditions (IC_90_, terminal time point; left) with doubling times for each condition in untreated Mtb (right). Hierarchical clustering of potencies as calculated with cosine distances and average linkage. IC_90_ is color scaled (log_10_ transformation) within each drug (Table S3). ND = Not determined. NA = not applicable. (B) Metrics from DiaMOND dose response curves. IC_50_ and IC_90_ are used to calculate drug interactions at the 50% and 90% levels of growth inhibition (FIC_50_ and FIC_90_, respectively). Three potency metrics are derived: AUC_25_ = normalized area under the curve until 25% inhibition, E_inf_ = theoretical maximum inhibition, and GR_inf_ = theoretical maximum normalized growth rate inhibition (Box and Materials and Methods). (C) Schematic data cube of the DiaMOND compendium. Mtb were treated with all 1-, 2-, and 3-way drug combinations (175 combinations) among 10-drugs in dose responses measured in 10-dose resolution in at least biological duplicate. Dose response measurements were made in eight *in vitro* models and at 3-4 time points, but we focus on 1-2 time points for analysis; therefore, this data cube represents ~25% of the total measurements made.

### Drug combination dose response measurements

For the DiaMOND compendium, we selected ten antibiotics in first- and second-line TB treatment regimens and for which there are abundant *in vivo* (mouse) data (Table 1, Table S2). These drugs include cell wall synthesis inhibitors (ethambutol, isoniazid, and pretomanid), rifamycin transcriptional inhibitors (rifampicin and rifapentine), protein synthesis inhibitor (linezolid), inhibitors of energy metabolism (bedaquiline and clofazimine), DNA replication inhibitor (moxifloxacin), and the antimycobacterial agent pyrazinamide (Fig. 1A, Table S2). We treated the Mtb Erdman strain carrying an autoluminescent reporter and measured both optical density (OD_600_) and luminescence at multiple time points after drug treatment. We observed a strong dependency in drug potency on *in vitro* model (Fig. 1A, inhibitory concentration to achieve 90% inhibition, IC_90_. Table S3), consistent with the idea that drug efficacy is influenced by bacterial stress *(44)*. We did not observe remarkable correlations in potency profiles by *in vitro* model. However, hierarchical clustering of drug potencies showed some groupings of drugs consistent with their target cell process (e.g., rifamycin transcriptional inhibitors group together, isoniazid and pretomanid - inhibitors of cell wall synthesis - group together). We also observed clustering of similar *in vitro* models. For example, potency profiles from growth media with short-chain fatty acids butyrate and valerate as the carbon source group together (Fig. 1A).

We observed condition-specific drug potencies consistent with previous reports, suggesting that the models we adapted for high-throughput drug response measurements may be predictive of outcomes in animals. For example, the activity of pyrazinamide in acidic and intracellular models and inactivity in the standard model (Table S2) was consistent with *in vitro (45, 46)* and animal studies *(47–49)*. We also observed pyrazinamide activity with lipid carbon sources, which has not been previously reported. As previously described, the rifamycins shared similar potency profiles with higher potency of rifapentine (Table S3) *(50)*. Bedaquiline was more potent in medium with lipids as the carbon source compared to standard medium with sugars as previously described *(51)*. Isoniazid potency was lower in the dormancy model, consistent with its inactivity towards non-replicating bacilli *(52–54)* and previous studies showing decreased efficacy in the presence of nitrite *(41)*. The wide range of single-drug responses and consistency with prior studies suggest that the *in vitro* models in this study produce non-redundant drug response data and form a validated set of conditions to model the lesion-specific variation in drug response.

Using these eight *in vitro* models, we constructed a compendium of systematic drug combination measurements by utilizing the DiaMOND method’s efficiency (Box). DiaMOND is a geometric optimization of the traditional checkerboard assay of drug-dose combinations. DiaMOND estimates the effect of combining drugs using a fraction of possible drug-dose combinations and focuses on the single drug and equipotent drug combination dose responses *(31)*. We measured all 1-, 2-, and 3-drug combination dose responses (totaling 175 combinations) in at least biological duplicate (Fig. 1B), resulting in a compendium of over 51,000 individual dose response curves. We focused our analysis on up to two timepoints per *in vitro* model to navigate this complex dataset. We chose the last time point (terminal, T) that is relative to the doubling rate (4 to 5 doublings for most models) and at a consistent treatment timepoint (constant, C) across *in vitro* models, ~7 days post treatment; Fig. 1A, Table S1. We also selected the measurement type that best benchmarks against colony forming units (OD_600_ for all models except intracellular and dormancy models, for which we used luminescence, Fig. S1). This selected dataset represents approximately one-quarter of the total number of compendium dose responses.

##### Box

DiaMOND (Diagonal measurement of n-way drug interactions) is a quantitative framework to efficiently measure drug interactions. The method is based on geometric sampling of traditional combination checkerboards and can be applied to any number of drugs in combination. Optical density (OD_600_) or luminescence measures are normalized to untreated controls and subtracted from 1 to obtain fractional growth inhibition. The concentrations to achieve a particular effect (e.g., concentration to achieve 90% growth inhibition, IC_90_, depicted in blue and orange circles) are experimentally determined for all single drugs so that dosing in subsequent measurement of drug combinations is equipotent (e.g., the IC_90_ should be dose #~7 for all drugs). Doses may be spaced linearly or logarithmically, but the spacing must be consistent between drugs. The single-drug dose responses (blue and orange boxes) and the equipotent drug combination dose response (black box) are highlighted. Drug interactions can be estimated using only the measurements from these boxes rather than the entire checkerboard by approximating the shape of isoboles (contours of equal effect). In the diagram, the isobole for IC_75_ is traced by the circles. If drug A and B are additive, the isobole would be a straight diagonal, and we calculate the expected IC_75_ on the combination dose response (orange square) where the dotted line intersects with the diagonal (combination) dose response curve. In this illustration, the combination reaches an IC_75_ at higher dose levels (orange circle) than the expected IC_75_, indicating an antagonistic interaction. The ratio of observed and expected doses (observed/ex-pected) is the fractional inhibitory concentration (FIC):

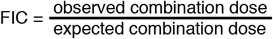

The DiaMOND methodology was used to obtain dose response data for every drug and drug combination measured over multiple time points. A Hill function was fit to these data and several potency and drug interactions metrics were derived from these dose response curves.

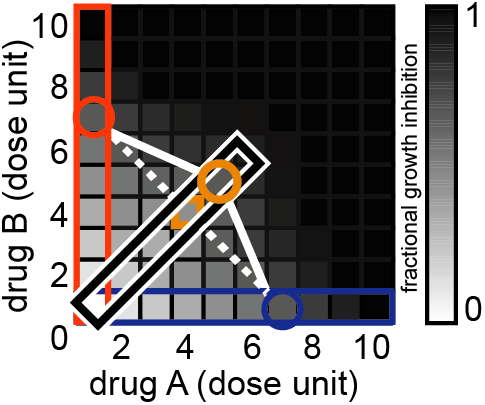

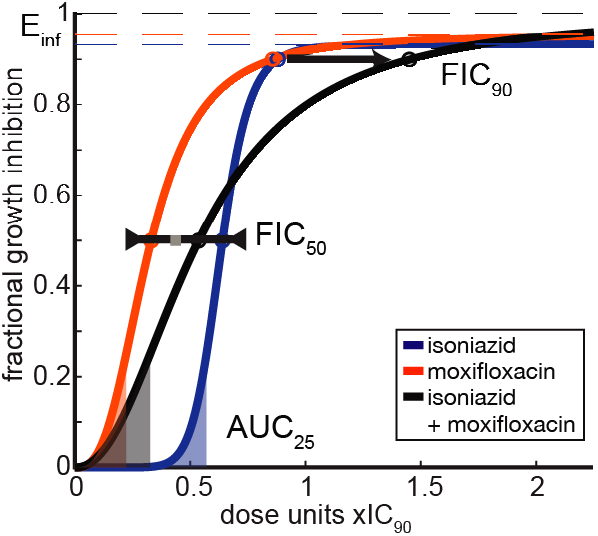

###### DiaMOND dose response metrics

**E_int_** (the maximum achievable effect): derived from the fitted Hill function (lower pane, dashed lines, colored by single drug or drug combination), E_inf_ describes the maximal achievable effect (upper asymptote, dashed lines) of a given drug or drug combination at a particular time point, where the maximum possible effect is 1.

**AUC_25_**: the area under the curve (AUC) simultaneously captures variation in potency and effect of a drug or drug combination, i.e., sensitivity to drug. AUC_25_ captures sensitivity to drug at concentrations with low growth inhibition. To compare low does potency to other drugs or drug combinations with different concentration ranges, we normalize the area by dividing by the IC_25_. The resulting AUC_25_values range from 0 (no effect) and 1 (potent).

FIC: drug interactions measure the effect of combining drugs on drug potency, i.e., the dose required to achieve a specific effect. The fractional inhibitory concentration (FIC) is the ratio of the observed combination dose (black circle) to achieve X effect over the expected combination dose (grey square), where FIC < 1 indicates synergy, FIC > 1 indicates antagonism, and FIC = 1 indicates additivity. In this example, the FIC_5g_ is approximately additive whereas the FIC_go_ is antagonistic, which is indicated by the relative position of the combination dose response (black) near (IC_5g_) and to the right (IC_90_) of the single dose response curves. We log transformation FICs to balance such that log_2_FIC < 0 is synergistic and log_2_FIC > 0 is antagonistic.

**GR_inf_**: Derived from the growth rate curve (not shown here, see (55) for details), GR_inf_ describes the maximal achievable effect of a drug or drug combination on the normalized growth rate, ranging between 1 and −1, where GR(c) is between 0 and 1 in the case of partial growth inhibition, GR(c) = 0 in the case of complete cytostasis, and GR < 0 indicates cell death. This unitless metric describes the effect of a drug on cells independent of doubling time, enabling comparison of drug effect on cells in different growth conditions.

We analyzed the single- and combination-drug treatments to derive potency and drug interaction information (see Box). With DiaMOND, we can quantify the degree and directionality of interactions at different growth inhibition levels using common null models (e.g., Loewe additivity and Bliss independence). Drug combinations that are more or less effective than expected based on single-drug behaviors are considered synergistic and antagonistic, respectively. Drug interactions are quantified with fractional inhibitory concentrations (FICs) at different growth inhibition levels (e.g., FIC_50_ and FIC_90_ are measured at the IC_50_ and IC_90_, respectively). FIC measurements were log-transformed to represent synergistic and antagonistic combinations with negative and positive log_2_(FIC) values, respectively. Drug interaction metrics based on Loewe additivity and Bliss independence were correlated (FIC_50_ and FIC_90_ for the constant and terminal time points, r=0.81, p < 2.2×10^−308^, Pearson’s correlation, Fig. S2); we, therefore, selected Loewe additivity as the null model for systematic analysis in the compendium (FIC_50_ and FIC_90_). Dose response curves provide treatment potency metrics at a low dose (AUC_25_; a normalized area under the curve to IC_25_, see Box) or high dose (E_if_ the maximum achievable effect). To compare potency across models where Mtb have different growth properties, we calculated the maximum achievable inhibition of normalized growth rate (GR_inf_; see Box), which allows direct comparison of treatment effects on cells with very different growth rates *(55)*. Though many other drug response metrics may be calculated from DiaMOND data, our analysis focused on these five metrics -- FIC_50_, FIC_90_, AUC_25_, E_inf_, GR_inf_ -- because they represent well-characterized and biologically interpretable aspects of drug interactions and potencies across low- and high-dose ranges.

### Drug synergy is uncommon and does not distinguish effective combinations

To identify patterns in drug interactions, we clustered the compendium drug interactions at the terminal time point in all eight growth environments, using 90% growth inhibition (log_2_(FIC_90_), Fig. 2A) and 50% growth inhibition (log_2_(FIC_50_), Fig. S3). Clustering did not reveal obvious model-wide synergy for any combination. Instead, we observed that most drug interactions were antagonistic (70% of FIC_90_>0), consistent with a general trend towards antagonism in drug interactions observed in other organisms *(31, 56–60)* and cancer *(61)*. The tendency towards antagonism depended on the growth model, with some conditions showing a balance between synergy and antagonism (intracellular and acidic), and others almost entirely antagonistic (cholesterol). Together these data suggest that synergy is a property of both drug and growth environment rather than an intrinsic property of the drug.

**Fig. 2.**
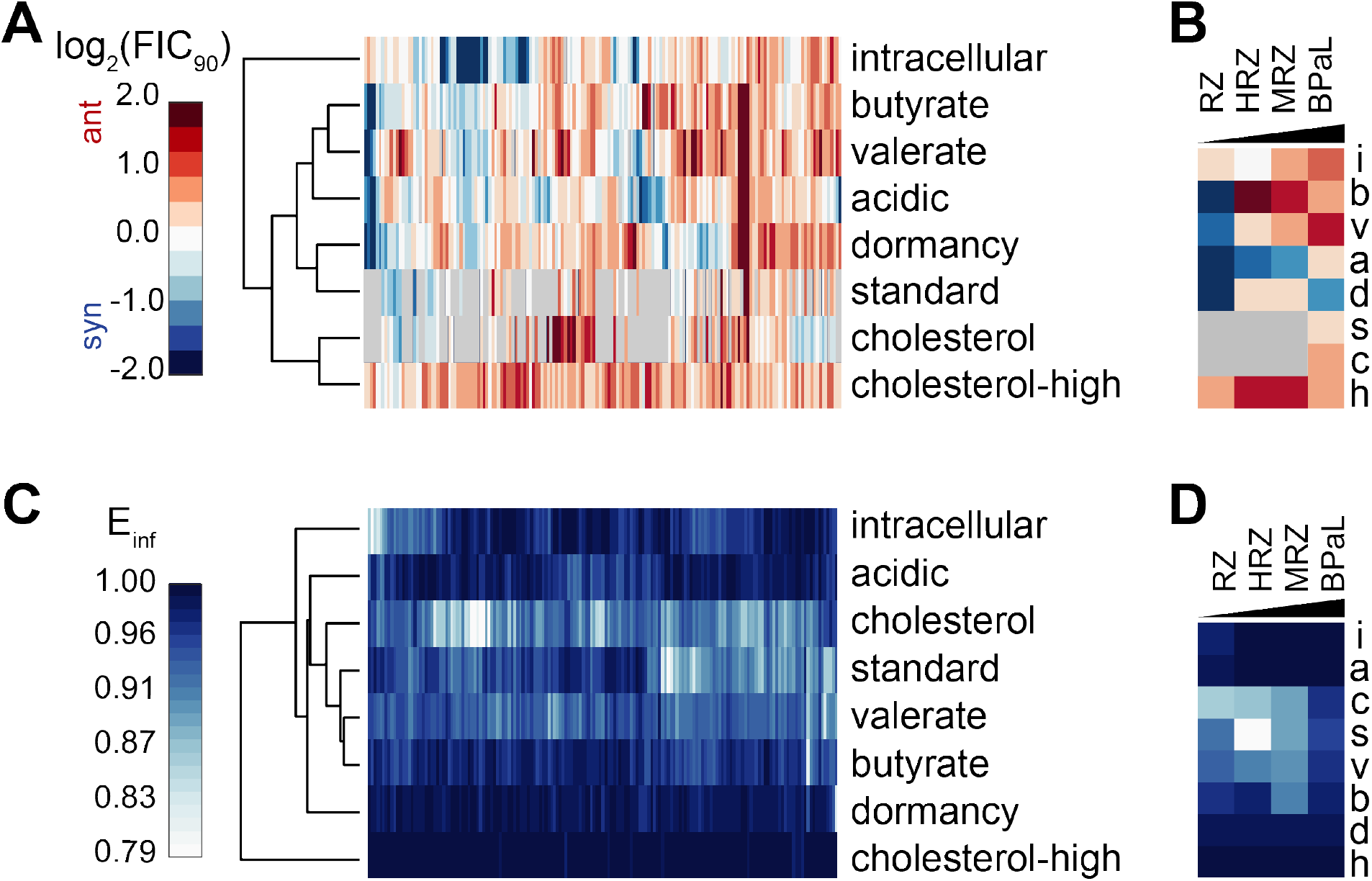
Drug interaction and potency patterns in the DiaMOND compendium. **(A)** Drug interaction profiles of all 2- and 3-drug combinations among the ten compendium drugs across the *in vitro* models (log_2_(FIC_90_) at the terminal time point, clustered based on cosine distance). **(B)** Drug interaction profiles of selected drug combinations ordered by mouse relapse outcome efficacy *(22, 64–71)*. See Table 1 for drug combination abbreviations. **(C)** Drug combination potency profiles of all 2- and 3-drug combinations among the ten compendium drugs across the *in vitro* models (E_inf_ at the terminal time point, clustered based on cosine distance). **(D)** Drug interaction profiles of selected drug combinations ordered by mouse relapse outcome efficacy *(22, 64–71)*. See Table 1 for drug combination abbreviations. gray = ND.

To understand whether combinations that tend toward *in vitro* synergy are more effective *in vivo*, we compared selected combinations with differences in disease relapse from the most commonly used mouse strains (e.g., BALB/c, C56BL/6, Swiss). The relapsing mouse models (RMM) evaluate drug efficacy months after cessation of drug treatment, somewhat analogous to the clinical measurement of relapse *(62, 63)*. We did not observe combination rank-ordering by synergy in any growth condition that matched efficacy in the RMM; e.g., BPaL>MRZ>HRZ>RZ (Fig. 2B) *(22, 64–71)*. Instead, we observed the 3-drug standard of care (HRZ) was the most synergistic drug combination, and BPaL was the most antagonistic among this subset (Fig. 2B). These examples suggest that drug interaction scores alone in the measured *in vitro* models were poor indicators of *in vivo* combination efficacy.

Synergistic drug combinations are not necessarily more effective than antagonistic combinations as the maximum effect of a combination can change independently of the drug interaction *(72)* (see Box). A tradeoff between synergy and efficacy appears to be important to consider when selecting effective drug combinations for treating other diseases (e.g., hepatitis C, HIV, and cancer), with maximum effect often being more important than synergy *(73, 74)*. To determine if the maximum effect could be used to prioritize combinations from the DiaMOND compendium, we clustered the E_inf_ (a measure of maximum dose response effect, see Box) for all compendium drug combinations in all eight *in vitro* models at the terminal time point (Fig. 2C). We observed a high maximum effect (E_inf_>0.9, Fig. 2C) in most combinations, consistent with the drugs’ known anti-Mtb effects. Dormancy and cholesterol-high models exhibited little variation in E_inf_, suggesting that neither condition had the dynamic range of maximum effect needed to discriminate among combinations or that all drug combinations are effective in these growth conditions for extended drug exposures. We compared E_inf_ profiles for the selected combinations we examined before, and we found that BPaL was more potent than HRZ or MRZ (Fig. 2D), consistent with animal outcomes of these regimens *(22, 64–71)*. These examples suggest that maximum achievable effect *in vitro* may be a stronger predictor of outcomes in mouse models than synergy. As with E_inf_, we observed correct rank ordering in some *in vitro* models by other potency metrics (AUC_25_ and GR_inf_) (Fig. S4), though we identified no drug combinations in the DiaMOND compendium that were maximally potent across all eight models (Fig. S5). The correct ordering of selected drug combinations by mouse outcome suggests that the DiaMOND compendium contains useful information for identifying efficacious drug combinations.

### DiaMOND metric signatures are predictive of treatment outcomes in the relapsing mouse model

We hypothesized that combinations of *in vitro* measurements could be compiled to model the *in vivo* microenvironments experienced by Mtb during drug treatment. We asked whether signatures of DiaMOND compendium measurements could distinguish drug combinations that were better than the standard of care in animal studies, HRZE or HRZ (Table 1). We classified 27 drug combinations that we measured in the compendium based on whether the treatment outcome in published RMM studies was better than the standard of care (C1) or not (C0) (Table S4). Principal component analysis (PCA) demonstrated that linear combinations of *in vitro* features could separate C0 and C1 drug combinations (Fig. 3A, Wilcoxon rank-sum, p<0.005, Table S5). Inspection of feature contributions to the principal component (PC) that best separates C0 and C1 drug combinations revealed many features related to cholesterol, standard, and valerate growth models (Fig. 3B). We also observed that potency metrics (AUC_25_, E_inf_, GR_inf_) are almost exclusively represented in the top 20 contributing features (Fig. 3B). Together these results suggest that effective separation of C1 and C0 drug combinations requires measurement of drug combination potency in multiple growth environments.

**Fig. 3.**
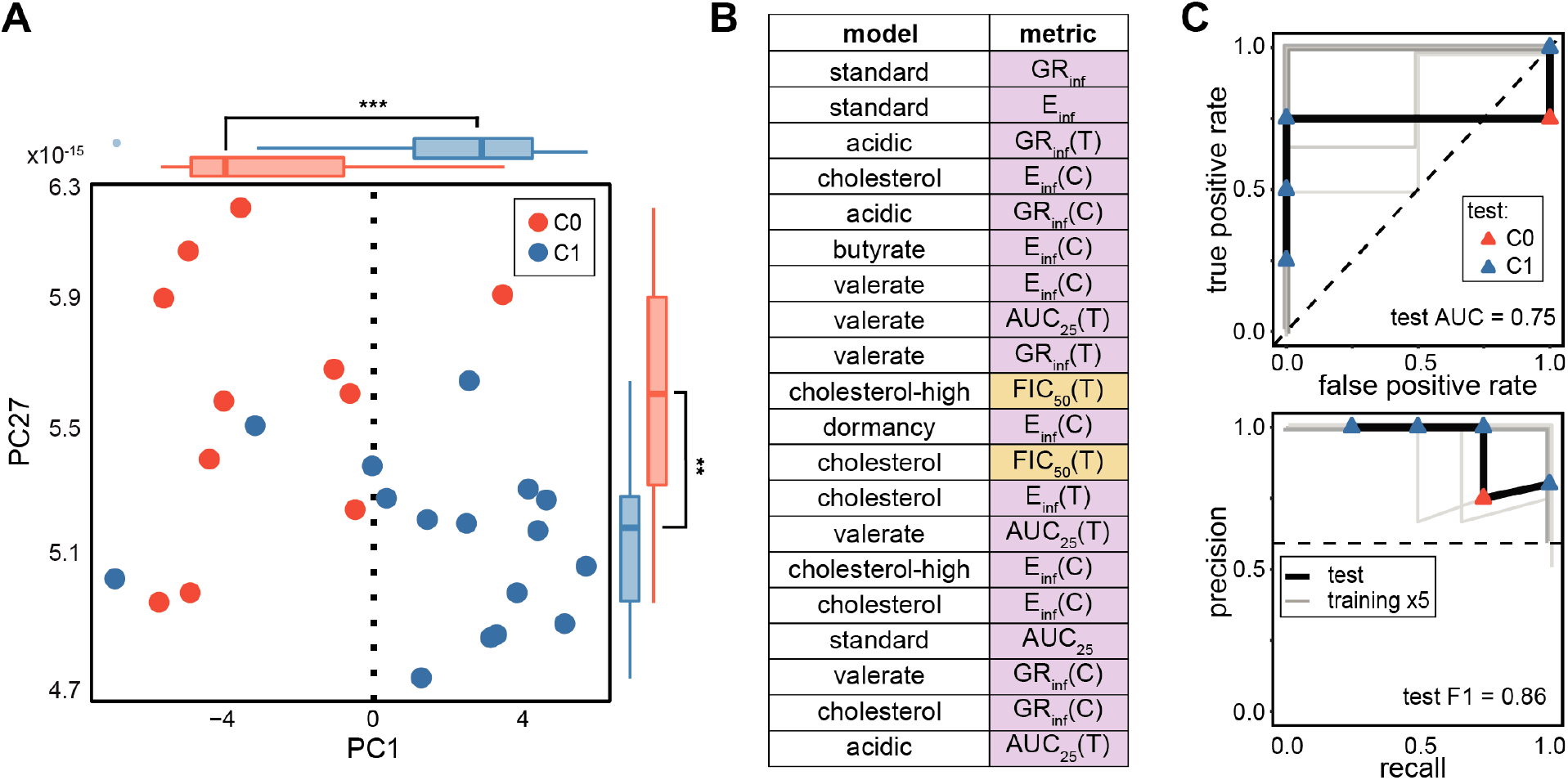
Prediction of combination treatment outcomes in the RMM with DiaMOND data. **(A)** PCA of DiaMOND data labeled by outcome in the RMM (C1 is better than the standard of care, blue; C0 is standard of care or worse, red) with the most discriminating two PCs shown. Outside the scatter plot are box and whisker plots of the distributions of C1 and C0 combinations along PC1 and PC27 (Wilcoxon rank-sum test: *** p<0.005. ** p<0.01). **(B)** Highest weighted features in PC1 with *in vitro* model (abbreviations in Fig. 1A) and metric type indicated. Metrics are classified and shaded according to whether they are related to drug combination potency (purple: AUC_25_, E_inf_, and GR_inf_) or drug interaction (orange: FIC_50_ and FIC_90_). **(C)** ROC curves (top panel, Table 1) and PR curves (bottom panel, Table 1) of a random forest-based classifier trained on all eight conditions in the DiaMOND compendium. The model is tested with high-order combinations (4- and 5-drug combinations) that were excluded from training. Training (gray lines each show one of five cross validations; lines are slightly offset to aid visualization) and test (black) performances are shown with lines. Test combinations are colored by outcome class as in (A). Performance metrics are shown on plots for test data (Area Under the ROC curve (AUC) and F1, harmonic mean of precision and recall, Table 1). Dashed lines indicate theoretical “no-skill” model performance.

To develop signatures of DiaMOND metrics that characterize C0 from C1 combinations, we trained binary classifiers with eight different machine learning (ML) methods to distinguish C0 and C1 drug combinations and compared their performance in 5-fold cross-validation (Table S6). We observed that nonlinear ensemble methods (Bayesian additive regression trees, random forest (RF), and gradient boosted trees) outperformed other ML algorithms, as measured by the area under the receiver operator characteristic (ROC) curve (AUC) and the F1 statistic, which is the harmonic mean of precision and recall (Table S6). We performed additional validation of the RF model by applying it to higher-order (4- and 5-way) drug combinations commonly used in preclinical and clinical tests that were not considered during model training (Table S7). The RF model accurately predicted outcomes (Fig. 3C, AUC=1, F1=0.86) and exhibited performance similar to what was estimated in cross-validation. Overall, the performance of ML models demonstrates that there is a strong signal in the DiaMOND compendium that is predictive of RMM drug combination efficacy.

We observed that some of the *in vitro* models in the DiaMOND compendium are well-represented among the top-ranked features in the classifying PCs (Fig. 3B). In contrast, other *in vitro* models are not present, suggesting that a subset of *in vitro* models may be sufficient to predict treatment outcome in the RMM. We asked whether classifiers using the DiaMOND compendium data from one *in vitro* model at a time were predictive of RMM outcome class. We observed that the data signal separating C0 and C1 drug combinations appeared in at least one PC for all eight *in vitro* models (Fig. 4A, Wilcoxon rank-sum test, p<0.05, Fig. S6). Furthermore, the five technically simpler models to work with exhibited clear C0 and C1 separation (Fig. 4A, *in vitro* models cholesterol, butyrate, standard, valerate, Fig. S6 *in vitro* model acidic).

**Fig. 4.**
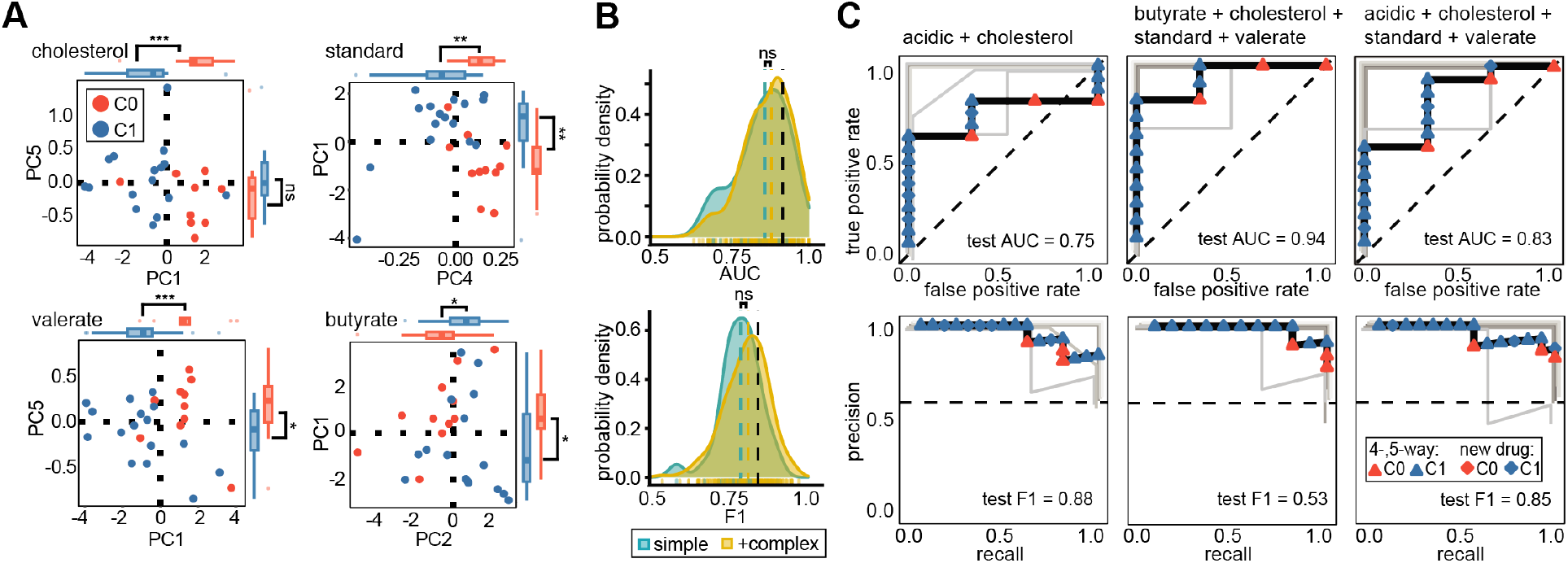
Prediction of combination therapy outcomes in the RMM using fewer *in vitro* models. **(A)** PCA plot of DiaMOND data labeled by outcome in the RMM (plots are labeled as in Fig. 3A). Each subplot is DiaMOND data from one *in vitro* condition plotted in the PC space with the most discriminating two PCs shown for each model. Outside the scatter plot are box and whisker plots of the distributions of C1 and C0 combinations along PC1 and PC2 (Wilcoxon rank test: *** p<0.005. ** p<0.01. * p<0.05. ns p>0.05). **(B)** Density distribution plots of estimated classifier performances from systematic survey of all possible *in vitro* model subsets. Distributions of ROC AUC (top) and F1 (bottom) are separated based on whether technically complex models (intracellular, cholesterol-high, dormancy) are included (yellow) or whether only simple conditions (acidic, butyrate, cholesterol, standard, valerate) are considered. Colored dashed lines indicate mean value for distribution. The estimated performances when using all *in vitro* models (as in Fig. 3) is shown with black dashed lines. Distributions are compared with a Wilcoxon rank sum test (ns = not significant). **(C)** Comparison of classification performances of three high-performance random forest classifiers using subsets of simple *in vitro* models. Training (gray lines each show one of five cross validations; lines are slightly offset when they are on top of each other) and test (black) performance is demonstrated with ROC (top) and PR (bottom) curves. Test combinations are colored by outcome class as in panel (A). Plot shapes indicate whether a test combination contained higher-order 4- and 5-drug combinations (triangle) or a combination containing a new drug (diamond-shape) not included in the compendium described in Fig. 1. Dashed lines indicate theoretical “noskill” model performance.

Though the single *in vitro* model classifiers were moderately predictive, they did not perform as well as the classifier trained using data from all eight *in vitro* models (Table S8). We asked whether another high-performing classifier could be derived using a subset of *in vitro* models. We systematically trained RF classifiers by considering all possible model combinations and observed that among the 255 possible combinations of *in vitro* models, 67 (26.3%) performed better than the classifier trained on all eight models. Furthermore, predictors including only the simpler *in vitro* models performed as well or better than those including the “complex” (intracellular, dormancy, cholesterol high) models (Fig. 4B, student’s t-test, p>0.05). We further validated the highest performing classifiers trained on the simple *in vitro* models by applying them to the higher-order (4- and 5-way) drug combinations as well as drug combinations involving antibiotics (delamanid, sutezolid, and SQ109, Table S1) that were not included in the compendium’s 10-drug set (Table S7). The high performance of classifiers on this validation set suggests that computationally combining simple *in vitro* models can produce classifiers that inform possible RMM outcomes (Fig. 4C). Additionally, the large number of classifiers that exhibit high accuracy and the shared DiaMOND compendium metrics among several *in vitro* models suggests that there may be multiple combinations of *in vitro* models that are predictive of outcomes in the RMM.

With many high performing RMM classifiers trained using subsets of the five simple *in vitro* models (Fig. 4C), we assessed whether the predicted RMM outcome for specific drug combinations would be consistent between these classifiers. The classifiers produce a probability that a drug combination belongs to each class (e.g., drug combination X belongs to C1 with 60% probability and C0 with 40% probability). The threshold probability is usually at 50% to assign the classification, but the probability can also rank combination classification likelihood. We tabulated the predicted probabilities of outcome for all combinations in the compendium, as well as the higher-order and new drug combination validation set, using the top-performing simple *in vitro* model classifiers shown in Fig. 4C. As we had previously observed, rank-ordering the percent probabilities within each classifier shows high predictive performance when evaluating the validation set. Among all predictions made for the compendium and validation combinations, we noted that 36% of drug combinations had discordant predictions among the three classifiers. We did not observe a consistent pattern in which a classifier was discordant. We next tested whether a consensus prediction could be generated by simply averaging the probabilities of the top three classifiers. We observed that the discordant combinations were clustered in the second quartile (probability of C1 around 25-50%), suggesting that classifiers are most prone to error for combinations that are C0. This may be due to the mild class imbalance in the training set (11 C0 and 16 C1 combinations). The consensus prediction was highly accurate (84% of validation set and 93% overall). Incorrect consensus predictions were at the border between C0 and C1 at 42-47% C1, indicating that the misclassification was due to ambiguity near the 50% decision boundary instead of strong classifier discordance. We conclude that a simple averaging of the probabilities generated by top classifiers is a practical means to construct an accurate consensus rank ordering for predicting drug combination response outcomes.

### DiaMOND metrics describe the efficacy of drug combination treatments in the C3HeB/FeJ mouse model

Given the success of ML classifiers to predict RMM outcomes, we next asked whether the DiaMOND compendium can be used to predict outcomes in other mouse models. Bactericidal activity in the most commonly used mouse strains (e.g., BALB/c, C56BL/6, Swiss) has been used extensively to evaluate drug combination effectiveness. Bactericidal activity in these models (bactericidal mouse model, BMM, Table 1) measures the reduction in bacterial burden by drug treatment immediately following drug treatment and can be assessed more quickly than relapse. Using the same analysis pipeline, we trained ML classifiers to recognize C0 or C1 drug combinations for the BMM outcome (Table S9) but observed that the classifier performance was only mildly predictive (AUC = 0.67, F1 = 0.40) (Fig. S7). Additional analysis of *in vitro* model subsets identified many predictors with improved performance, but this improvement did not generalize to test data. Moderate model training performance and poor generalizability to new data suggest that the drug combination information needed for BMM outcome predictions may be difficult to capture with the *in vitro* models developed and used in this study.

The C3HeB/FeJ (HeB) mouse strain has become important for TB regimen development because the disease pathology is more similar to humans than other mouse strains *(26, 27, 75)*. This includes the formation of caseous, necrotic granulomas that are characterized by low oxygen content (hypoxia) *(27, 75, 76)* and differential drug penetrance *(77, 78)*. These lesions also contain large numbers of extracellular, non-replicating bacteria *(24, 76)*. Like other mouse studies, those with HeB mice use bactericidal (BHeB) and relapse outcomes to determine drug effectiveness. Fewer drug combinations have been tested and published using HeB mice than other mouse strains. The DiaMOND compendium contained too few measured combinations to train ML classifiers. When we integrated the compendium combinations with higher-order drug combinations, we obtained a total of 16 combinations (Table S10) for the BHeB outcome, which was sufficient to train ML classifiers. However, we were not able to do the same for the relapse outcome, where we had four total combinations, even after augmenting with higher-order information.

To understand if DiaMOND metrics distinguish C0 and C1 BHeB combinations using this expanded training dataset, we evaluated class separation with PCA. We observed significant separation of BHeB outcome classes along the third PC (PC3) (Fig. 5A, p<0.005, Table S11). We then examined the top 10 features in PC3 by contribution (Fig. 5B) and found that the *in vitro* models and metrics were distinct from those we observed in the RMM analysis (Fig. 3B). Notably, the metrics for the BHeB were entirely drug interactions (Fig. 5B), and the presence of the dormancy model in the top ten features was of particular interest because we expected hypoxia-induced dormancy to be a microenvironment specific to the C3HeB/FeJ mice *(27, 75, 76)*. Using the same approach described for RMM, we developed accurate RF models to classify BHeB C1 and C0 combinations (Fig 5C, all in vitro models, AUC=0.9, F1=0.80). Systematic evaluation of RF classifiers using all possible combinations of *in vitro* model subsets revealed that complex models did not improve performance (Fig. S8). Specifically, we found that models without dormancy perform as well as those with it (Fig. S9). As with the RMM classifiers, we identified *in vitro* model subsets that performed better than all models together trained for the BHeB outcome (37 (12.9%)). Lipid and acidic *in vitro* models featured prominently among the most accurate classifiers. Together, these analyses demonstrate that the DiaMOND compendium data predicts outcomes in two pathologically distinct mouse models, suggesting that enough key information can be captured by simple *in vitro* models to prioritize combination therapies for animal model tests.

**Fig. 5.**
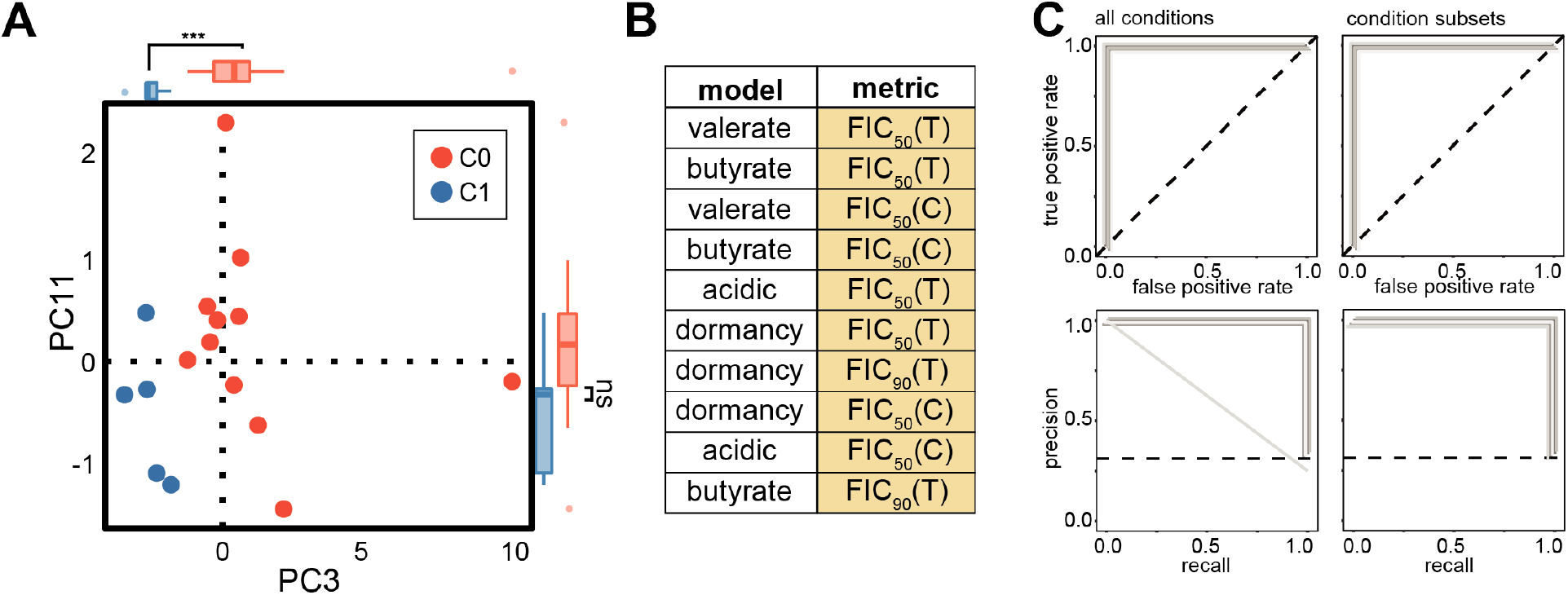
Signatures of DiaMOND data to describe outcome in the C3HeB/FeJ (BHeB) mouse model. **(A)** PCA plot of DiaMOND data labeled by outcome in the BHeB (plot labels are as in Fig. 3A). **(B)** Highest weighted features in PC3 with *in vitro* model and metric type indicated (features are as described in Fig. 3B). **(C)** Machine learning performance plots for training with 5-fold cross validation (each in a gray line) with ROC (top) and PR (bottom) curves for models trained using all eight conditions (left) and three high-performing subsets of conditions (right: acidic + butyrate + valerate, cholesterol + valerate, and standard + valerate). Subsets had perfect training performance (AUC = 1.0, Table S7). Dashed lines indicate theoretical “no-skill” model performance.

### Potency and antagonism are correlated with improved outcomes in mouse models

The signatures of DiaMOND data describing outcomes in RMM and BHeB highlighted that potency metrics were key predictors for RMM, while drug interactions were key for BHeB outcome classification. To understand whether C0 and C1 drug combinations showed significant differences in these metrics, we examined the top five features from the most discriminatory PCs for both mouse models. Univariate analysis revealed significant differences between three of the top five features for the RMM outcome (Fig. 6A. Wilcoxon rank-sum, p<0.05). In each of these five features, potency was higher in C1 combinations than C0 combinations, which is consistent with expectations of increased potency for the most effective drug combinations. The top features describing BHeB outcomes are drug interactions (Fig. 6B), and three of the top five exhibited significant differences between C0 and C1 combinations (Wilcoxon rank-sum, p<0.05). For all five drug interaction features, C1 was the more antagonistic class. That antagonistic drug combinations may be more favorable is consistent with the results of our comparison of BPaL to the standard of care (HRZ, Fig. 2B). We found that different metric types (potency or interactions) may provide information that maps to different outcome types (bactericidal or relapse) in animal studies. Furthermore, our analysis suggests that high potency and antagonism in *in vitro* assays may be characteristics of favorable drug combinations.

**Fig. 6.**
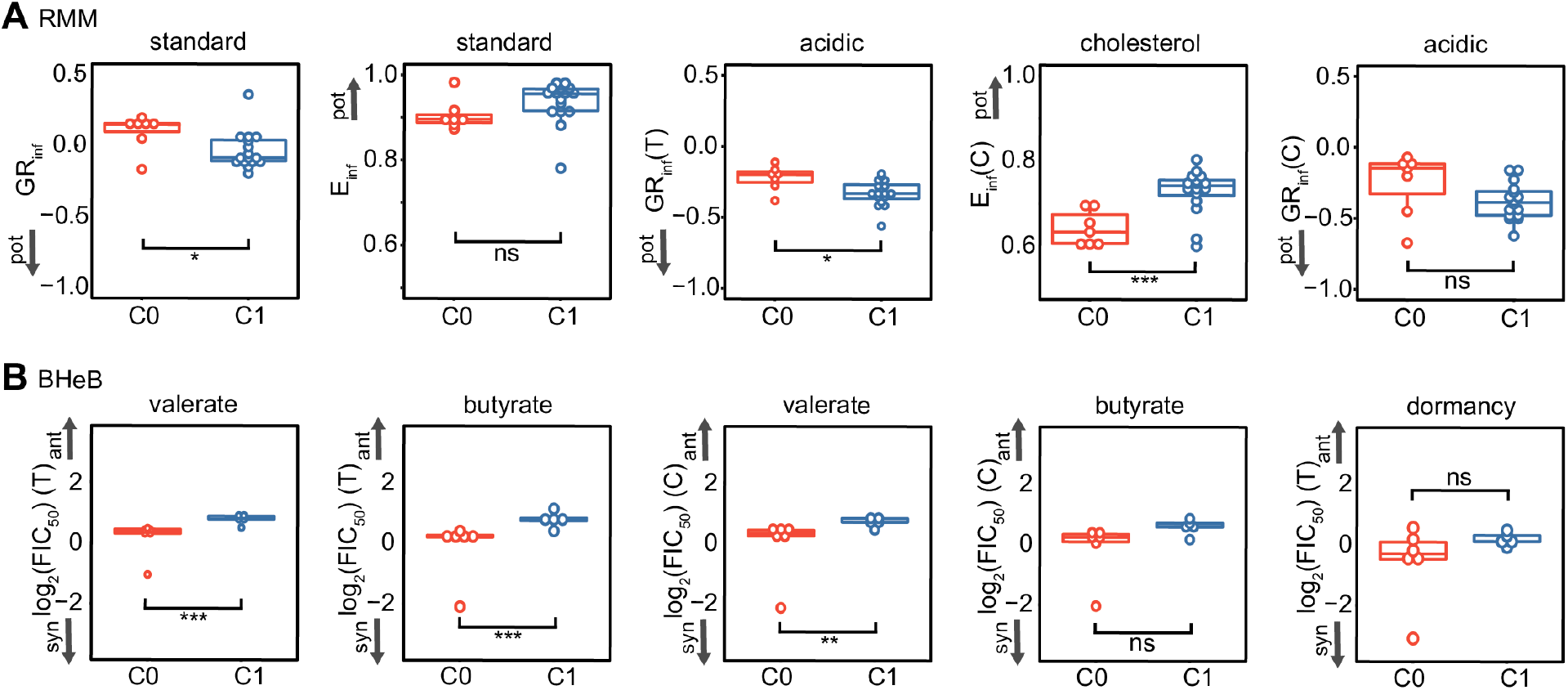
Properties of signature potency and drug interaction characteristics that describe RMM and BHeB combination treatment outcomes. Values of the five highest weighted features in the most discriminatory PC are compared for C1 and C0 combinations in the RMM **(A)** and BHeB models **(B)** using dot and box plots. The top features in RMM are potency metrics whereas the top features are drug interaction metrics in BHeB. High vs. low potency (pot) and synergy (syn) vs. antagonism (ant) is indicated with arrows on each subplot. (Wilcoxon rank test: *** p<0.005. ** p<0.01. * p<0.05. ns p>0.05).

## Discussion

Our goal in this study was to develop a pipeline to efficiently prioritize drug combinations early in the TB regimen design process. Most *in vitro* drug efficacy studies utilize single growth conditions, which have not been clearly mapped to *in vivo* outcomes *(24, 30)*. Furthermore, conflicting results from multiple *in vitro* models have not been readily resolvable. We hypothesized that treatment efficacy *in vivo* could be modeled as a “sum-of-parts” of the complex microenvironment. Therefore, we generated a dataset that profiles drug combination effects against Mtb in eight different *in vitro* growth environments. With this comprehensive drug combination data compendium, we identified signatures of potencies and drug interactions in specific *in vitro* models that distinguish whether drug combinations are better than the standard of care in two important preclinical mouse models. We found that ML classifiers were accurate predictors of mouse disease relapse using data from only a few simple *in vitro* models. These classifiers were validated with higher-order (4- and 5-drugs) combinations and had predictive power for combinations with drugs not included in the model training. Together, our study establishes a practical approach to prioritize combination therapies using economical, scalable, and expandable *in vitro* measurements.

Synergy is often assumed to be a property of optimized combination therapies because synergistic drugs are more effective together than expected based on single-drug efficacies alone. Our mapping of the DiaMOND compendium onto outcomes in two different mouse models challenges this notion. In the relapsing mouse model, drug interactions were not key features for classification; instead, the potency measures from the drug dose response curves were the most important predictors of outcome (Fig. 3B). Our findings are consistent with reports of treatment in hepatitis C, cancer, and HIV *(73, 74, 79)* that show a tradeoff between maximizing synergy and potency of a drug combination. Maximizing potency was often more important than synergy in treating these diseases with multidrug therapies *(73, 74, 79)*. Antagonism was prevalent in our compendium (Fig. 2A), and we found that antagonism was characteristic of more efficacious drug combinations for the C3HeB/FeJ bactericidal model (Fig. 6B, C1 more antagonistic than C0). Partnering the most potent drugs together during regimen design may be generating highly potent combinations but biasing these combinations towards antagonistic drug interactions. Bedaquiline, pretomanid, and linezolid were recently found to be more potent in treating mice infected with the Mtb HN878 strain than the H37Rv strain *(80)*. When combined, the drugs antagonized each other for treating Mtb strain HN878-infected mice. Despite this antagonism, the BPaL combination was highly effective at curing mice infected with either Mtb strain. These *in vivo* results are consistent with our findings that BPaL is a highly potent but antagonistic drug combination for *in vitro* treatment of Mtb Erdman. One view of how drugs in combination exert their effect on cell populations is that each drug targets a different subpopulation rather than multiple drugs targeting the same cells *(73, 79)*. Drug interactions would then explain how well a drug acts on the cellular population that was not susceptible to the other drugs in the multidrug treatment. This leads to the hypothesis that very potent drugs that alone can kill most of the cells in a population would achieve high maximum effect when combined but may tend toward antagonism rather than synergy. Study of the multidrug anticancer therapy R-CHOP (Table 1) supports this hypothesis *(73)*, and an expanded study using more antibiotics could be used to test this hypothesis in tuberculosis. Our study suggests that for TB, potent drug combinations should be prioritized for further study and should not necessarily be deprioritized if they are antagonistic in *in vitro* assays.

Our approach enabled us to determine the relative importance of specific *in vitro* models to predict outcomes in mice, thereby serving to validate which growth conditions map to *in vivo* responses. We note that signatures including several *in vitro* models perform better than signatures using data from only one *in vitro* model, perhaps because the lesion microenvironment is complex and constitutes multiple stressors that affect Mtb drug response. It may also indicate that none of the physiological states imposed by the *in vitro* models we used in this study was so dominant that it drives Mtb *in vivo* drug treatment response. We chose several lipids to serve as carbon sources in our *in vitro* models because of the important role of lipids and cholesterol specifically for Mtb growth, survival, and infectivity *(81–90)*. The cholesterol *in vitro* model was the top-performing single *in vitro* model classifier for the RMM outcome and performed almost as well as the classifier with all *in vitro* models. This is consistent with the importance of cholesterol metabolism for Mtb survival and infectivity *(24, 85–87)*. We also observed that other lipid-rich environments (modeled by using short-chain fatty acids butyrate or valerate as carbon sources) induced distinct drug response patterns and that the best classifiers for both RMM and BHeB outcomes utilized metrics from multiple lipid-rich growth conditions. These findings suggest that measuring drug combination responses with a suite of simple growth environments may be sufficient to model the complex lipid environment encountered in TB lesions.

Mtb in the RMM mouse strains are thought to be primarily intracellular *(24)*, and intracellular Mtb are exposed to the acidification of the phagolysosome *(36, 37)*. Therefore, we expected the acidic growth environment to be a driver in classifiers for the RMM. We found that measurements from the acidic growth environment alone were not strongly predictive of outcomes in the RMM but that these metrics were prominent in the best mixed-condition classifiers. Furthermore, other single growth environment models perform better than the acidic model (Table S6). These results indicate that response to acidic stress is important for Mtb intracellular survival to drug treatment in the RMM, but adaptation to other environmental factors (such as lipid carbon sources) are important drivers of treatment response. We also observed that the acidic model was prominent among the best classifiers for the bactericidal outcome in the C3HeB/FeJ mouse strain (BHeB outcome). The C3HeB/FeJ mice are noted for the formation of the caseous necrotic granulomas (type II lesions, *(76)*) that have been shown to have a neutral pH (pH>7) *(91)* and with primarily extracellular Mtb *(76)*. However, these animals have abundant intracellular bacteria in other lesion types and within macrophages that acidify the intracellular Mtb compartments *(77)*, which may explain why acidic growth environments are important predictors of drug response in this mouse model.

The microenvironments in TB granulomas are complex, yet we were able to combine measurement in a “sum-of-parts” approach from relatively simple growth environments to model treatment outcomes (Fig. 4B, Figure S8). These results indicate that there is predictive drug combination response information obtained from simple *in vitro* models that only needs to be combined correctly to predict drug treatment outcomes in mice. The practical implication is that researchers can choose a subset of the most amenable *in vitro* models for performing drug combination experiments and still retain predictive capacity. We successfully model RMM and BHeB outcomes using this compendium but not the BMM outcome, suggesting that there may be other important factors or emergent properties of complex environments encountered during infection that were not included in our measurement set. These factors may include other carbon sources, nutrient availability, iron limitation, oxygen tension, human serum, or other entry mechanisms into dormancy (e.g., via different lipids or other combinations of stressors) *(91–93)*. Macrophage activation status has also been linked to Mtb drug susceptibility *(94)*, and therefore models of cytokine-induced activation of macrophages may be more relevant to specific *in vivo* outcomes. Future studies measuring drug combination response in other and more complex *in vitro* models may permit accurate modeling of the BMM outcome and improve the accuracy of predictions of outcomes in RMM, BHeB.

We anticipate that our *in vitro*-to-*in vivo* pipeline for drug combination predictions may be applied to study treatment outcomes in other animals and clinical studies. Predictions using our RMM classifiers suggest the potential for using DiaMOND data to model responses in the clinic. The moxifloxacin containing regimens, HRZM and MRZE, were expected to shorten treatment time in humans by two months based on preclinical mouse studies but did not show non-inferiority to HRZE in the ReMOX clinical trial *(95)*. A meta-analysis of mouse relapse outcome studies after the trial completion revealed that the treatment shortening of these combinations was expected to be smaller than initially thought and that perhaps could explain the ReMOX trial results *(62)*. A recent PET/CT study in humans of 14-day treatment regimens also found MRZE and HRZE to be comparable in the reduction of lesion volume *(96)*. The consensus RMM classifier predictions from our study predicted that the MRZE and HRZM combinations were not likely to show treatment improvement (47% and 53% chance of being C1, respectively), consistent with the ReMOX trial outcome, PET/CT study, and mouse meta-analysis. Based on our prediction probabilities, MRZE and HRZM would not have been prioritized for further study compared to other combinations, like BPaMZ (83% chance of being C1), that are being evaluated in ongoing clinical trials *(97)*. Another example of the potential utility of the DiaMOND compendium for clinical predictions comes from the “Study 31” clinical trial. The preliminary results from “Study 31” show treatment shortening of the continuation phase using PHZM compared with the standard treatment of HRZE (intensive) followed by HR (continuation) *(9)*. A third treatment arm (PHZE) did not show improvement compared with the standard treatment. Similar to our ReMOX predictions, our consensus prediction indicated PHZE to have a low probability for treatment improvement over standard of care (42% chance of being C1) despite the mouse outcome indicating C1 classification. The ReMOX and “Study 31” examples suggest that the DiaMOND compendium contains information that is relevant to understanding clinical outcomes while contradictory to the mouse outcomes. Together these results indicate that the DiaMOND compendium could be used in future modeling and clinical outcomes predictions *(98, 99)*.

Several changes to the experimental design may improve this pipeline. The importance of potency metrics in signatures of combination efficacy is perhaps surprising given that we design combination dose responses to have equipotent combinations of each drug. There is growing evidence that there is differential drug penetration into the lesions where Mtb is found *(78)*, which would lead to non-equipotent levels of drug reaching Mtb cells. Utilizing pharmacokinetic data to design drug combinations may increase this approach’s utility and power and lead to a more predictive DiaMOND compendium dataset. The current standard of care and other new regimens (e.g., “Study 31” and SimpliciTB) involve intensive and continuation phases of treatment. Including sequential treatments in an experimental approach could help understand how prior treatment sensitizes the bacterial population to future treatment regimens. One reason to use combination therapy for TB is to slow the acquisition of drug resistance. A systematic study of the drug combination space in different growth environments can also be used to investigate the evolution of drug resistance. For example, antagonistic drug interactions have been shown to suppress the evolution of drug resistance *(56, 73, 100, 101)*, and the evolution of drug resistance can be tied to the growth rate and duration of drug exposure *(56, 80, 102, 103)*. Finally, the depth of the DiaMOND compendium may be well-complemented with transcriptomic data of drug response to prioritize drug combinations based on predicted mechanisms of drug interaction *(104)*.

TB is not the only disease that benefits from combination therapy. We expect that our pipeline may be adapted and applied to optimize multidrug regimens for other diseases, including cancers, HIV/AIDs, and multi-drug resistant bacterial pathogens. Beyond our use of the TB DiaMOND compendium to describe combination efficacies in mouse models, we anticipate that this dataset may be used for other systematic studies of drug combination response.

## Materials and Methods

### Strains and media

*M. tuberculosis* Erdman strain was transformed with pMV306hsp+LuxG13 to generate an autoluminescent strain that was used for all experiments in this study (Addgene plasmid # 26161; http://n2t.net/addgene:26161; RRID:Addgene_26161) *(105)*, see Supplemental Materials). We used the mouse cell line, J774 as a model of intracellular residency because J774 cells have been used as a macrophage-like cell line to study early infection processes and Mtb drug response to complex host-like intracellular environment *(48, 106)*.

Standard 7H9 Middlebrook medium supplemented with 0.2% glycerol, 10% OADC (0.5g/L oleic acid, 50g/L albumin, 20g/L dextrose and 0.04g/L catalase) and 0.05% Tween-80 with 25 μg/mL kanamycin was used for Mtb strain maintenance. Growth and culturing were performed at 37°C with aeration unless noted. All *in vitro* model media were buffered with 100 mM 3-(N-morpholino)propanesulfonic acid (MOPS, pH 7), unless noted, and filter-sterilized prior to use. The acidic model was based on the standard 7H9 Middlebrook media above and buffered with 100 mM 2-(N-morpholino)ethanesulfonic acid (MES) to pH 5.7. For acclimation to lipid carbon sources, a base medium consisting of 7H9 powder (4.7g/L), fatty acid-free BSA (0.5g/L), NaCl (100mM) and tyloxapol (0.05%) with 25 μg/mL kanamycin was used and the lipids sodium butyrate (5mM, final concentration), valeric acid (0.1% final concentration) or cholesterol (0.05mM or 0.2mM final concentration) were added to the base medium. For the cholesterol media, a cholesterol stock solution (100mM) was first prepared by dissolving cholesterol in a 1:1 (v/v) mixture of ethanol and tyloxapol and heated to 80°C for 30 minutes and added to pre-warmed (37°C) base medium *(32)*. The dormancy media was based on the butyrate media with the addition of sodium nitrate (5mM) as a terminal electron acceptor *(38, 41–43)* J774 cells were cultured as previously described *(106)*. Briefly, J774 cells were cultured in high glucose DMEM supplemented with 2mM L-glutamine, 1mM sodium pyruvate, and 10% heat-inactivated fetal bovine serum (FBS) at 37°C in 5% CO2. Media was changed every one-three days and cells passaged at ~80% confluence. Standard 7H10 Middlebrook agar plates supplemented with 0.5% glycerol, 10% OADC, 0.05% Tween-80 and 25 μg/mL kanamycin were used for enumerating colonies.

### Mtb in vitro model acclimation

Mtb were inoculated into standard 7H9 Middlebrook medium, grown to mid-log phase (optical density, OD_600_ ~0.5-0.7) and were subcultured for less than two weeks prior to acclimation to assay medium. For acclimation to standard and acidic media, Mtb cells were diluted into assay media at a starting density of OD_600_ = 0.05, acclimated for 3-5 doubling times or until they reached mid-log phase (OD_600_ ~0.5-0.7), diluted to OD_600_ = 0.05 and grown back to mid-log phase before use in DiaMOND assays.

Similar to standard, and acidic conditions, Mtb were acclimated to butyrate, and valerate media and acclimated cells were frozen for use in assays. Frozen acclimated Mtb in butyrate and valerate media were inoculated into assay media, grown to mid-log phase (OD_600_ ~0.5-0.7), diluted into fresh lipid media at a starting concentration of OD_600_ = 0.05 and grown back to mid-log phase (OD_600_ ~0.5-0.7) and used for DiaMOND assays. The dormancy model used Mtb acclimated to butyrate medium grown to mid-log phase (OD_600_ ~0.5-0.7) and then diluted to a starting OD_600_ 0.05 in dormancy media. For the dormancy model (d), cells were incubated at 37°C without aeration for 28 days, which reduced autoluminescence close to media-only background levels, which we interpret as being dormant with very low metabolic activity.

Mtb growth on cholesterol media slowed without the exchange of fresh medium. Cholesterol and cholesterol-high acclimation were similar to standard and acidic conditions with fresh media exchanges every seven days to ensure continued growth. Mtb acclimated between 14 and 28 days were used for assays. Mtb growth rate on cholesterol-high was faster (four day doubling time) than cholesterol (seven day doubling time).

For the intracellular model, J774 cells were plated at 375,000 cells/mL in 384-well plates and cultured overnight, expecting ~one doubling prior to infection. Mtb grown to mid-log phase in standard media was syringe-passed 8 times with a 25-gauge needle to reach a single-cell suspension, and J774s were infected with Mtb at MOI 2 for 24 hours followed by drug treatment for 5 days.

### Drugs, dose responses, and dispensing

The drugs used in this study are listed in Table 1. All drugs were reconstituted and diluted in DMSO except for PZA for the intracellular model; to avoid exceeding the DMSO limit (0.5%) in the intracellular condition, PZA was diluted in 1x phosphate-buffered saline with 0.01% Triton-X. Drugs were dispensed with an HP D300e digital dispenser, and locations were randomized to reduce plate effects. For each *in vitro* model, the concentration to achieve 90% inhibition (IC_90_) was determined. IC_90_ were used to design combination dose responses with equipotent mixtures of drugs *(31)*. A ten-dose resolution with 1.5- or 2-fold dose spacing was used for all experiments.

### Treatment and DiaMOND assays

Mtb were acclimated to *in vitro* model media prior to drug treatment as described above. For acidic, butyrate, cholesterol, cholesterol-high, standard, and valerate models: 50μL of acclimated Mtb at the indicated density was added to each well in 384-well plates containing freshly dispensed drugs and incubated at 37°C in humidified bags to prevent evaporation. Edge wells contained media but were not used for assays. For the dormancy model: Mtb were acclimated as described above, gently resuspended, and 20μL of dormant Mtb culture was transferred to each well on the assay plates. Plates were sealed with PCR seals to reduce oxygen exposure during drug treatment and incubated for seven days. We measured regrowth after drug treatment as a readout of drug effect during dormancy. Therefore, after drug treatment, plate seals were removed, 80μL of standard media was added to each well, and plates were incubated at 37°C in humidified bags to prevent evaporation. For the intracellular model: drugs were printed into media-only plates and transferred onto infected J774 cells 24 hours after Mtb infection. To accommodate quality control assessment, we included multiple untreated and positive drug treatment controls in each plate as well as uninfected J774 cells for the intracellular model. (See Supplementary Materials).

### Plate measurements

Luminescence and OD_600_ measurements were made at 3-5 time points per sample on a Synergy Neo2 Hybrid Multi-Mode Reader. Time points were based on the approximate doubling time of each model. To simplify the analysis, we generally compare time points at either a relatively similar time point (constant) or time ~4-5x doubling times after drug exposure (terminal time point). Constant and terminal time points correspond to the same set of measurements for the standard and intracellular *in vitro* models (constant/terminal, CT). For the dormancy model, plate readings were made during recovery in standard media, and time points were selected based on doubling time in standard media. For the dormancy and intracellular models, OD_600_ measurements could not reflect Mtb biomass alone, so only luminescence measurements are used. Autoluminescence has been demonstrated as a proxy for Mtb cell growth *(105)* and viability in response to drug treatment *(107, 108)*. To benchmark changes in luminescence to changes in growth in our conditions, we performed a series of drug treatment experiments in the dormancy and intracellular models (see Supplementary Materials and Fig. S1). Briefly, cells were treated as described above, followed by plating treated cells on 7H10 plates to enumerate colony forming units (CFU). Portions of the luminescence dose response curve that correlated with CFU changes were considered indicative of growth inhibition, and metrics derived from these portions of the curve were used for analysis.

### Data processing and metric calculation

Data processing and dose response metric calculation were performed using custom MATLAB scripts. In brief, raw data were background-subtracted using the median of media-only wells and normalized to the mean of untreated wells within each plate. For the intracellular model, uninfected macrophages provided the background (rather than media only) for subtraction from raw data, and subsequently, data was normalized to (infected) untreated within each plate. A 3-parameter Hill function was fit to each dose response (single drug or combination). Inhibitory concentrations (ICs) were calculated based on the Hill curve parameters. The area under the curve at 25% inhibition (AUC_25_) was calculated using the integral of the fit curves from 0 to the 25% inhibitory concentration (IC_25_) and normalized to the IC_25_, allowing comparisons between drug combinations. Drug interaction scores were quantified by the fractional inhibitory concentration (FIC) using Loewe additivity and Bliss independence (See Box). FICs calculated by Loewe additivity and Bliss independence were well correlated, and neither model was observed to suffer from significant bias relative to the order of the drug combination *(109)* (Supplementary Material); therefore, we proceeded to analyze drug interactions based on Loewe additivity. The growth rate inhibition (GR) metrics were calculated as previously described *(55)*. See Supplementary Materials for details on data processing and analysis.

### Data quality

Experiments were performed in a minimum of biological triplicate. Comparisons of data between plates and between experimental days required data quality control assessment. Each dose response was assigned a quality score that takes into account the overall quality of the data from a plate, the quality of fit of the Hill function, the single drug dose responsiveness from an experiment, and in the case of drug combinations, the equipotency in the drug combination dose responses (Supplementary Materials). In brief, plate data quality was assessed with a Z’-score using multiple untreated (negative) and complete inhibition treatment (positive) wells in each plate. The fitting of the Hill function was assessed by the coefficient of determination (R^2^) of the fit as well as the closeness of the E_inf_ for each fit to the maximum observed effect for each dose response curve. Drug combination equipotency was assessed by comparing the proportional combinations normalized to their respective MICs and the idealized combination of drugs if they were perfectly equipotent. Dose responses with poor quality scores were excluded from further analysis.

### Computational Analyses

Biological replicate dose response and drug interaction data passing quality control were averaged. Means of replicate data were used for all downstream analyses unless noted. Hierarchical clustering was performed using cosine distance, and heatmaps with complete linkage dendrograms were generated using MATLAB. Other data preparation and visualizations were performed in R studio (version 3.5.3) using the tidyverse environment packages and ggplot2 and ggpubr packages for visualization. Data table import and export were performed in R using the openxl and readxls packages

PCA was performed in R using the prcomp function from the stats package with each feature scaled to have unit variance before PCA. Some features were missing data; e.g., FIC_90_ metrics were missing because single drugs did not achieve IC_90_. Features with more than 35% missing data points were excluded from PCA. The remaining missing values were imputed using the mean of the corresponding input features (mean imputation) *(110)*.

### Machine learning

The machine learning in R (mlr v2.17.0) package was used for all machine learning tasks involving projections of the original features onto the principal component (PC) space as the input features and classified drug combination outcome (C0 or C1) as labels.

### Feature selection, feature number optimization, and model validation

The Kruskal-Wallis test was used to rank order the PC input features for ML based on the ability to discriminate outcome classes C0 and C1. As there were a limited number of drug combinations, we aimed to reduce the number of features used in the model. A Monte-Carlo resampling strategy was used to split the training data into 70/30% training/test partitions, to which we applied grid search to find the number of features that produced the largest test AUC. This feature number optimization was repeated five times for each training set, and the smallest feature set from the five iterations was chosen as the final training feature set. Models were trained on the full set of training data, and performance on new data was estimated using standard 5-fold cross validation. Validation was performed by projecting new data onto the PC space used for the model training and testing model classification performance.

### Machine learner packages

Upon feature selection, machine learning algorithms were compared using standard 5-fold cross validation. The performance was evaluated using the AUC and the F-score (F1). The mlr package made possible on-demand loading of learners from other R packages, including Bayesian additive regression tree (bartMachine, v1.2.5.1), random forest (randomForestSRC, v2.9.3), extreme gradient boosting (xgboost, v1.1.1.1), logistic regression (stats), naive bayes (e1071, v1.7-3), support vector machine (e1071, v1.7-3), and weighted k-nearest neighbors (kknn, v1.3.1).

### Statistical Analysis

Differences between outcome class groups for DiaMOND features or PCs were assessed by means (IC_90_ averages), medians (class comparisons), and standard deviation of drug combinations from each outcome group in each *in vitro* model. Because data normality could not easily be assessed with small numbers of drug combinations in each group, the Wilcoxon rank-sum test was used to compare outcome group means for statistical significance. Student’s t-tests were used for testing hypotheses of differences between model performance distributions. The hypothesis that Loewe and Bliss interaction (FIC) scores were correlated was tested using Pearson correlations. Statistical analyses were performed using the stats, ggpubr, or the rstatrix packages in R version 3.5.3.

## Supplementary Materials

Materials and Methods

Fig. S1. Benchmarking cell viability with luminescence measurements.

Fig. S2. Comparing null reference models for drug interaction scoring.

Fig. S3. Intermediate potency drug interaction profiles.

Fig. S4. Alternative potency metric profiles for selected drug combinations.

Fig. S5. Alternative potency metric profiles for DiaMOND compendium.

Fig. S6. Outcome class separation in single *in vitro* model principal component analyses (PCAs).

Fig. S7. BMM classifier performance on training and test data.

Fig. S8. BHeB *in vitro* model subset model performance distributions.

Fig. S9. BHeB *in vitro* model with and without dormancy model performance distributions.

Table S1. Experiment time points and estimated growth amounts for *in vitro* models.

Table S2. Drug information table.

Table S3. Drug IC_90_ for *in vitro* models.

Table S4. Drug combinations with RMM outcomes for 2- and 3-way drug combinations.

Table S5. RMM PC class separation.

Table S6. Machine learning algorithm benchmarking performance metrics.

Table S7. RMM model validation set.

Table S8. RMM single *in vitro* model classifier performance.

Table S9. Drug combinations with BMM outcomes for 1-, 2- and 3-way drug combinations.

Table S10. Drug combinations with BHeB outcomes for 1-, 2- and 3-way drug combinations.

Table S11. BHeB PC class separation.

## Acknowledgments

We thank members of the Aldridge laboratory, V. Dartois, R. Isberg, K. Mdluli, J. Mecsas, A. Palmer, J. Silverman, and S. Tan for insightful discussion. Bedaquiline and pretomanid were provided by the NIH AIDS Reagent Program and the TB Alliance, respectively. pMV306hsp+LuxG13 was a gift from Brian Robertson and Siouxsie Wiles.

## Author contributions

JLF, TG, NV, YD, and BBA conceived and designed the experiments. JLF, TG, NV, and YD performed the experiments. JLF, TG, NV, YD, MO, AS, and BBA conceived and designed the computational analysis. JLF, TG, NV, YD, and MO performed the computational analysis. The manuscript was written by JLF and BBA. All authors contributed to interpretation of the results and editing of the manuscript.

