## Supplemental Materials for "Systematic measurement of combination drug landscapes to predict *in vivo* treatment outcomes for tuberculosis"

**Supplementary Materials:**

Materials and Methods

*Generation of autoluminescent Mtb strain*

Autoluminescent *M. tuberculosis* strain was generated by transforming the Erdman parent strain with pMV306hsp+LuxG13, resulting in a single copy chromosomal integration of the bacterial luciferase operon. The pMV306hsp+LuxG13 plasmid contains a reorganized and codon-optimized bacterial luciferase operon for maximum mycobacterial light production (*105*). Briefly, the construct was electroporated into Mtb, and kanamycin resistant colonies were isolated and tested for auto-luminescence. These positive strains were expanded in standard 7H9 supplemented media and frozen down. The frozen stocks were used as the starting strains for *in vitro* model acclimation and drug combination experiments.

*Dose centering*

For every *in vitro* model, each single drug was tested to identify the $IC_{90}$ (concentration to inhibit 90% growth). Each dose response was ten units and the $IC_{90}$ for single drugs was designed to be between dose 6 and dose 9. Drug combinations were designed for dosing to be equipotent around the $IC_{90,}$ and doses were spaced 1.5x or 2x apart to capture the drug's full range of response.

*Benchmarking luminescence measurements*

Decreases in autoluminescent Mtb have been shown to correspond to decreases in optical density and colony forming units (*105, 108*), specifically in response to drug treatment (*107, 108*). Luminescence must be used for the intracellular model because optical density measures both the mammalian and Mtb cells. We chose to use luminescence measurements for the dormancy model because the optical density measurements were highly variable. To more directly compare luminescence dose responsiveness from the intracellular and dormancy models to the optical density dose responsiveness of the other *in vitro* models, we sought to benchmark luminescence to growth inhibition. In the intracellular model, drug treatment was performed as described in Material and Methods. Luminescence was measured 6 days after infection (5 days after addition of drugs, Constant/Terminal time point). Mtb were then lysed from macrophages with 0.01% sodium dodecyl sulfate (SDS) in distilled water for 15 minutes at 37°C, 10-fold serially diluted with standard 7H9 media and plated on 7H10 Middlebrook agar for colony forming unit (CFU) enumeration. Mtb dormancy was established as described in Materials and Methods and treated with drugs. At the appropriate constant and terminal time point, luminescence was measured. Mtb were then 10-fold serially diluted with standard 7H9 media and plated on 7H10 Middlebrook agar for CFU enumeration. Normalized luminescence inhibition was calculated as described below and correlation assessed using "polyfit" in MATLAB.

*Data analysis*

Data quality and processing

Several measures were taken to ensure high quality measurements for this dataset. Every plate contained untreated bacteria and in-plate standards (specified drug at the $IC_{90}$ as determined from dose centering) as follows: for butyrate, cholesterol, acidic, standard, and valerate conditions: isoniazid and linezolid; for cholesterol-high: isoniazid; for intracellular and dormancy: moxifloxacin. Each dose response curve was assigned a quality control score that took into account the quality of growth and drug treatment within a plate (Z' score), the success of capturing the dose response range (dose space score), the quality of the fit of the Hill function to the data ($E_{inf}$ score and $R^2$ score), and the equipotency of drug combinations (angle score).

A Z' score was calculated for every plate to measure separation of strong positives from untreated in each experiment (See Z' calculation section). If the Z' score of a plate was less than 0.3, the plate was assigned a score of 2. If the Z' score was between 0.3 and 0.5 the plate score was 1. Plates with Z' score greater than 0.5, indicating that there was moderate discriminatory power between untreated and maximum treated wells, were assigned a plate score of 0.

To assess the quality of dose response range for a drug or combination, the number of data points collected for that drug/combination that fell between 10% and 90% inhibition was quantified. If this number was greater than or equal to 3, the "dose space score" assigned was 0. If the number was 0, the dose space score assigned was 2. If the number fell between 0 and 3, the dose space score assigned was 1.

To assess the quality of fit of the Hill function, two measures were quantified: an "$E_{inf}$ score" and the "$R^2$ score." The $E_{inf}$ score assessed how the $E_{inf}$ compared to the effect at the highest tested dose of a drug or combination ($E_{max}$). If the absolute value difference between the $E_{inf}$ and $E_{max}$ is greater than 0.1, the assigned $E_{inf}$ score is 2. If the absolute value difference is between 0.05 and 0.1, the assigned $E_{inf}$ score is 1. Below 0.05 was assigned 0. For each fit, an associated $R^2$ was also calculated. If the $R^2$ was $< 0.7$, the $R^2$ score assigned was 2. If the $R^2$ fell between 0.7 and 0.9, the $R^2$ score was 1. Greater than 0.9 was assigned 0.

To assess the equipotency of drugs in combination dose responses (an important consideration for DiaMOND calculations), an "angle score" was calculated for combinations. This score measured the difference between the true diagonal measured (the combination of N drugs in an N-way combination) and the ideal diagonal if every drug in that combination were precisely centered around the $IC_{90}$. If the difference between the angles (in degrees) was greater than 22.5, the angle score was 2. Between 10 and 22.5 received a score of 1, and less than 10 received a score of 0. All these scores were combined to compute a "composite score" for every single drug and combination. For single drugs, the composite score was calculated by:

$$Composite\ score\ (single) = \frac{1}{3}(Plate\ Score) + \frac{1}{3}(Dose\ Space\ Score) + \frac{1}{6}\left(E_{inf}\ Score + \ R^2\ Score\right)$$

For drug combinations, the composite score took into account the data quality of the underlying singles and the combination itself, where the underlying single score for each single drug in an N-way combination was calculated by:

$$Underlying\ single\ score = \frac{1}{2}(Plate\ Score) + \frac{1}{4}\left(E_{inf}\ Score + \ R^2\ Score\right)$$

and the resulting combination score was calculated by:

$$\begin{aligned} &Composite\ score\ (combination) \\ &\quad = \frac{2}{3}\left(\frac{1}{3}Plate\ Score + \frac{1}{3}E_{inf}\ Score + \frac{1}{6}(R^2\ Score + Angle\ Score)\right) \\ &\quad + \frac{1}{3*N}\sum Underlying\ single\ scores \end{aligned}$$

The composite score ranges between 0 and 2, where 0 is optimal and 2 is poor. Drugs or combinations with a composite score greater than or equal to 1 were rejected for further analysis. In addition, all fit Hill functions, and raw data for all single drugs in every experiment were checked manually; drugs that behaved unexpectedly or where the $IC_{90}$ was below dose 5, at or above dose 10 of the dose response were removed along with all combinations that contained that drug.

Z' calculation

To determine which conditions showed reproducible drug responses, we calculated a Z' score. A Z' score was calculated by the formula

$$Z' = 1 - \frac{3(\sigma_{pos} + \sigma_{neg})}{|\mu_{pos} - \mu_{neg}|}$$

where σ is the standard deviation and μ is the mean of the positive (pos) and negative (neg) controls, respectively, of those populations. We used the Z' to assess *in vitro* model reproducibility and for in-plate quality control.

<u>Fitting Hill function to dose response data</u>

We used a 3-parameter Hill function where for any concentration *x* of a drug or drug combination, *Hill(x)* describes the effect at that concentration as defined

$$Hill(x) = \frac{E_{inf}}{1 + \left(\frac{EC50}{x}\right)^{h}}$$

where $E_{inf}$ describes the maximum effect achievable by a given drug or drug combination, $EC_{50}$ describes the concentration to achieve 50% of the maximum effect, and h is the Hill slope. Data was normalized to untreated, and therefore the bottom asymptote of the Hill function was bound at 0. We found that dose response data had non-constant error variance in the media-based growth conditions, and therefore we implemented weights when we fit the Hill function to our data such that

$$Weights\ for\ Hill(x_i) = \frac{1}{stdev(growth\ measurement\ for\ biological\ replicates\ of\ dose\ i)}$$

Data points with lower variance are assigned more weight than samples with high variance when fitting. Two fitting algorithms were used: the Levenberg-Marquardt algorithm and the trust-region-reflective algorithm, each with different constraints as permitted by each algorithm. The

Levenberg-Marquardt algorithm does not allow bound constraints while trust-region-reflective does; therefore, we restrict the $E_{inf}$ to not go above 1 with the trust-region-reflective solution but cannot apply that bound to the Levenberg-Marquardt solutions. This occasionally results in a fit from Levenberg-Marquardt where the $E_{inf}$ is much greater than 1 (i.e., 100% inhibition). As this has no biological meaning, such fits are not appropriate for our purposes, and we discarded those fits. To assess fit quality, an $R^2$ was calculated; the fit from the two algorithms with the higher $R^2$ was chosen.

Occasionally, the $E_{inf}$ of the fit Hill functions was far above or below the maximum measured effect ($E_{max}$). We categorized these dose responses into those that had a maximum effect asymptote in the normalized data or those that had no clear asymptote. Accurate representation of maximum achievable effect was important for our analysis. Therefore, we attempted to improve agreement between the $E_{inf}$ of the fitted Hill function with the $E_{max}$ using a custom refitting strategy. For original fits that had an $E_{inf}$ below the $E_{max}$, the refitting of the Hill function had the lower bound of $E_{inf}$ parameter space constrained to within 1.25% of the $E_{max}$. For fits that had an $E_{inf}$ above the $E_{max}$, the refitting of the Hill function had the upper bound of $E_{inf}$ parameter space constrained to within 1.25% of the $E_{max}$. Refitting with and without weights were assessed using $R^2$ values. Refits with the highest $R^2$ were chosen as the final fit for a given dose response curve. Additionally, the Hill coefficient during fitting had an upper bound at 10.

For all fits, the inhibitory concentration (IC) to achieve 10, 25, 50, 75, and 90% growth inhibition or kill was calculated according to the formula:

$$IC_N(drug\ x) = \frac{EC50_{drug\ x}}{\sqrt[h_{drug\ x}]{\frac{E_{inf,drug\ x}}{inhibition\ level} - 1}}$$

The area under the curve at 25% inhibition (a measure of low potency) was calculated:

$$AUC25 = \frac{\int_0^{IC25} Hill(x)\,dx}{IC25}$$

<u>Drug interaction quantification</u>

DiaMOND is a tool that employs geometric optimization of the combination dose space to quantify drug interactions with fewer measurements(*31*). Drug interactions are quantified by the fractional inhibitory concentration (FIC) score, which is the ratio of the observed combination dose to achieve a certain effect over the expected combination dose to achieve that same effect. An FIC < 1 is considered synergistic, FIC > 1 is antagonistic, and FIC = 1 indicates additivity. For this study, we calculated FIC scores at various growth inhibition levels. The expected dose is based on the behavior of the single drugs in the combination. We employed two null models to calculate the expected combination dose: Loewe (dose) additivity and Bliss independence(*111*). Loewe additivity assumes dose additivity; that is, the effect of drugs in combination is determined by the sum of their normalized doses. By the Loewe model, the expected combination dose to achieve any inhibition level falls on the hyperplane defined by the single doses of each drug to achieve that inhibition level. The intersection of the combination dose line and the hyperplane is the expected combination dose. Bliss independence assumes response additivity; that is, drugs in combination act independently such that one cannot interfere with another. By the Bliss model, the effect of drugs in combination can be predicted by multiplying the effects of the singles, and thus the expected combination dose to achieve any inhibition level can be calculated.

*Growth rate (GR) metrics*

In addition to growth inhibition at static time points, growth rate inhibition was calculated as described previously (*55*). Normalized growth rate inhibition is calculated according to the formula:

$$GR(c) = 2^{\frac{log_2\left(\frac{x(c)}{x_0}\right)}{log_2\left(\frac{x_{unt}}{x_0}\right)}}$$

where x(c) is the $OD_{600}$ or luminescence readout for a given drug at concentration c at a specific time point, $x_0$ is the $OD_{600}$ or luminescence readout at time point 0 (T0), and $x_{unt}$ is the $OD_{600}$ or luminescence readout of the untreated population at the same specific time point. This was computed for all the concentrations in each dose response curve and then a 3-parameter Hill function was fit to the data using the trust-region-reflective algorithm:

$$GR(c) = GR_{inf} + \frac{1 - GR_{inf}}{1 + \left(\frac{c}{EC50_{GR}}\right)^{h_{GR}}}$$

where $GR_{inf}$ is the maximum growth rate inhibition achievable by a given drug or combination, $EC50_{GR}$ is the dose to achieve 50% of the maximum growth rate inhibition, and $h_{GR}$ is the Hill slope of the curve.

Fig. S1.

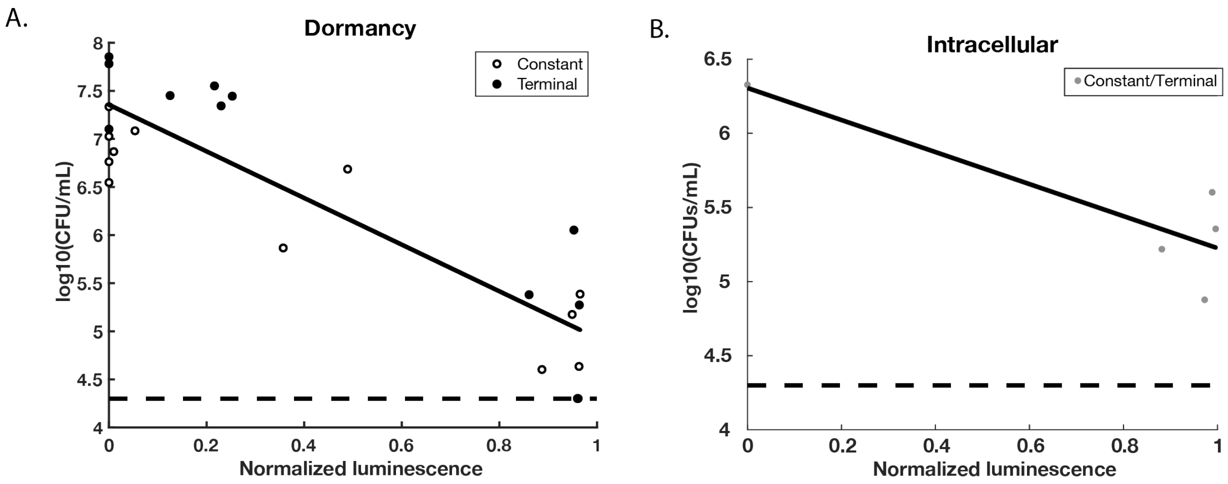

**Fig. S1.** Benchmarking cell viability with luminescence measurements. Normalized luminescence inhibition is compared with the resulting CFU/mL for **(A)** dormancy or **(B)** intracellular *in vitro* conditions. Cells were untreated or treated with drugs as in main experiments. At indicated times (**(A)** constant or terminal, or **(B)** constant/terminal), luminescence was measured, cells were removed from multiwell plates, diluted and CFU enumerated. Luminescence was normalized to untreated as described in Methods and Materials. Dashed line indicates the limit of detection for CFU/mL. Solid line indicates linear regression line ((A) r = -0.89, p-value = $8.9 \times 10^{-9}$, (B) r = -0.86, p-value = 0.06, using Pearson correlation).

Fig. S2.

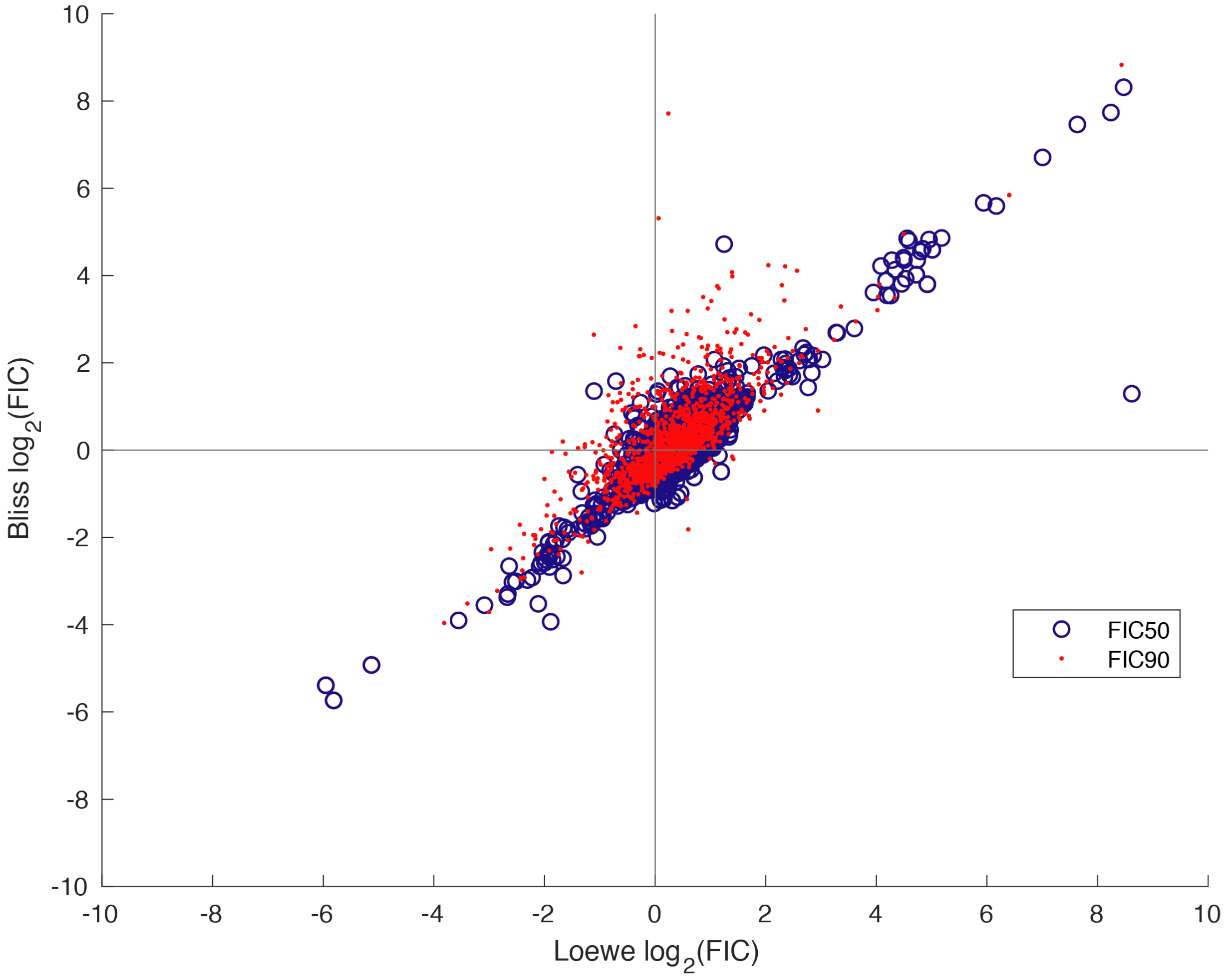

**Fig. S2.** Comparing null reference models for drug interaction scoring. The $\log_2(FIC_{50})$ and $\log_2(FIC_{90})$ scores calculated using either the Bliss independence or the Loewe additivity null models for each DiaMOND compendium 2- and 3-drug combination at the constant and terminal time points are compared using Pearson correlation, $r=0.81$, $p < 2.2x10^{-308}$).

Fig. S3.

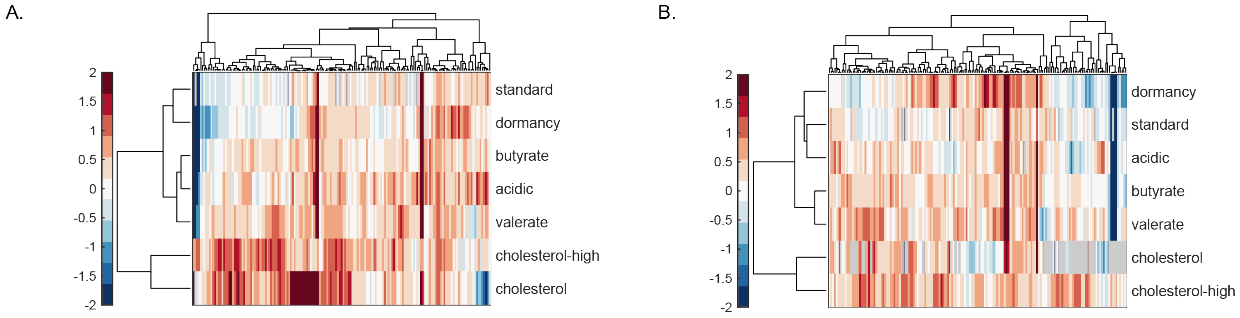

**Fig. S3.** Intermediate potency drug interaction profiles. Drug interaction profiles of each DiaMOND compendium 2- and 3-drug combination among the 10-drugs in the compendium across the *in vitro* models ($\log_2(FIC_{50})$) at the **(A)** terminal time point, and **(B)** the constant time point. Clustered based on cosine distance with complete linkage as described in Figure 2.

Fig. S4.

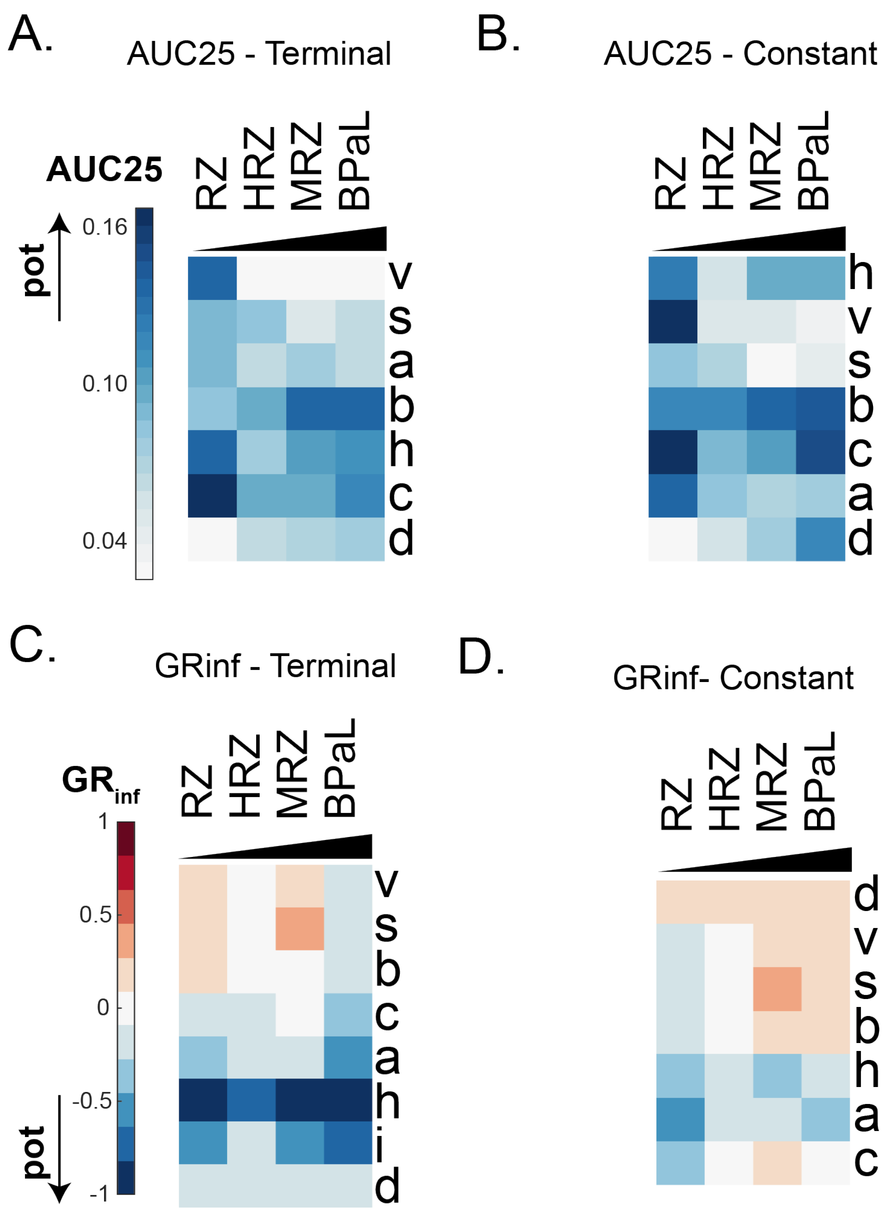

**Fig. S4.** Profiles of selected drug combinations with mouse relapse outcomes across the *in vitro* models for $AUC_{25}$ at the **(A)** terminal and **(B)** constant time points and $GR_{inf}$ at the **(C)** terminal and **(D)** constant time points. Drug combinations are ordered by relapse outcome efficacy as in Figure 2 (RZ is least and BPaL is most effective in group)

Fig. S5.

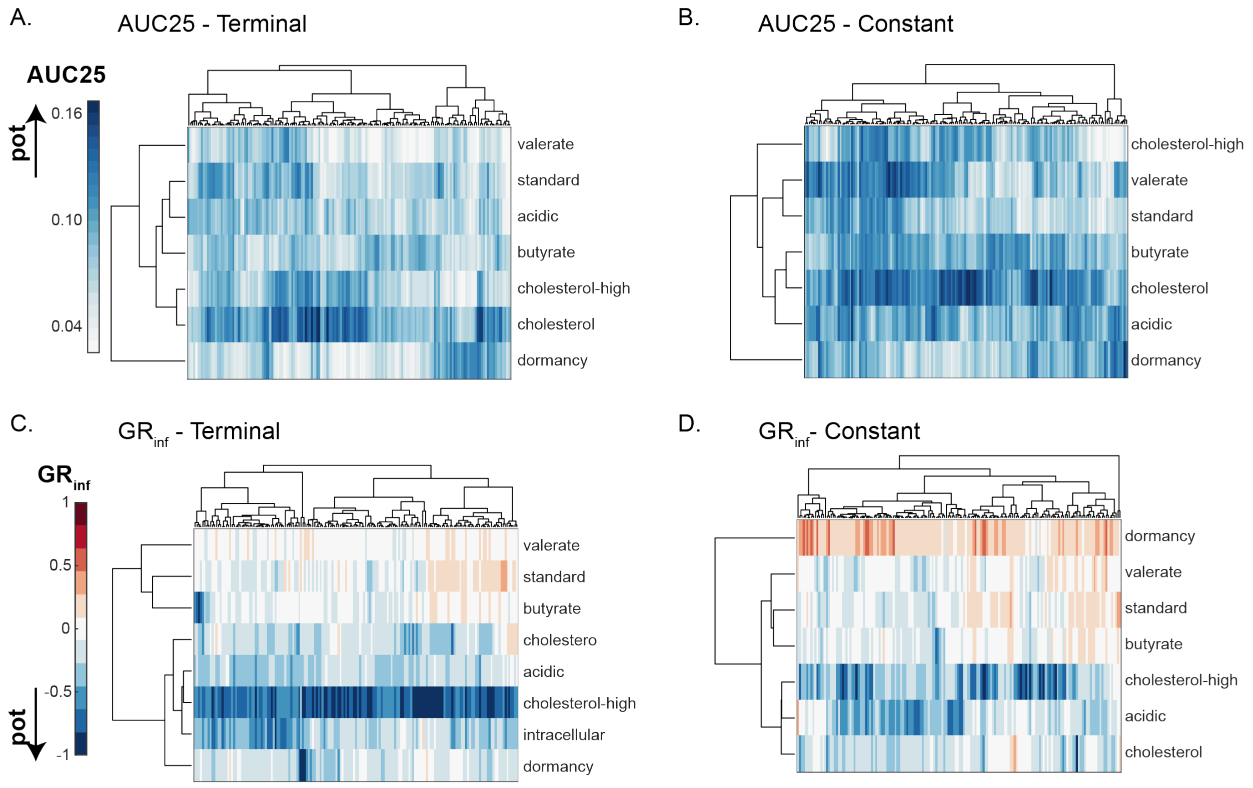

**Fig. S5.** Alternative potency metric profiles for DiaMOND compendium. Profiles of each DiaMOND compendium 2- and 3-drug combination among the 10-drugs in the compendium across the *in vitro* models for $AUC_{25}$ at the **(A)** terminal and **(B)** constant time points and $GR_{inf}$ at the **(C)** terminal and **(D)** constant time points. Clustered based on cosine distance with complete linkage as described in Figure 2.

Fig. S6.

A.

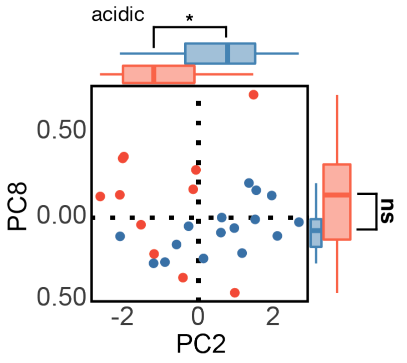

B.

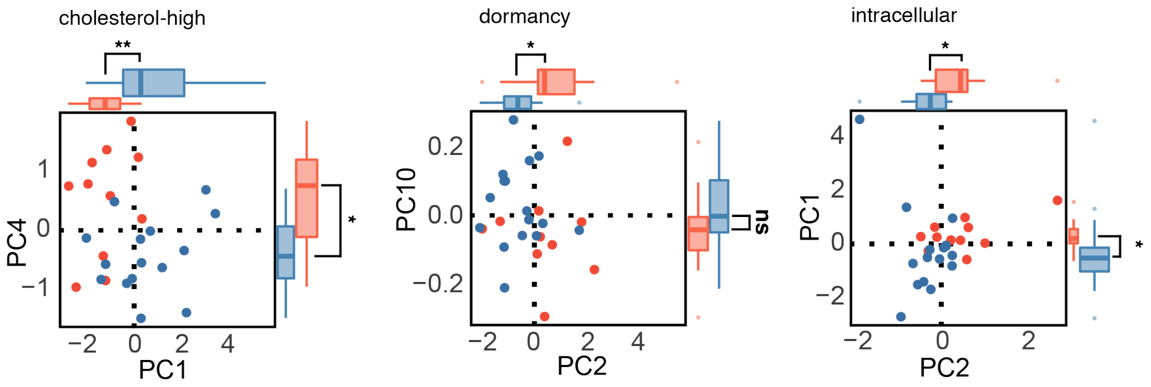

**Fig. S6.** Outcome class separation in single *in vitro* model principal component analyses (PCAs). PCA of DiaMOND compendium data for drug combinations with mouse relapse outcomes were performed, and top separating principal components (PCs) are plotted. Data are labeled by mouse relapsing mouse model (RMM) outcome (plots are labeled as in Fig. 3A). The *in vitro* models are separated into logistically **(A)** simple (acidic) or **(B)** complex (intracellular, cholesterol-high, dormancy). Outside the scatter plot are box and whisker plots of the distributions of C1 and C0 combinations along the indicated PCs (Wilcoxon rank test: *** p<0.005. ** p<0.01. * p<0.05).

Fig. S7.

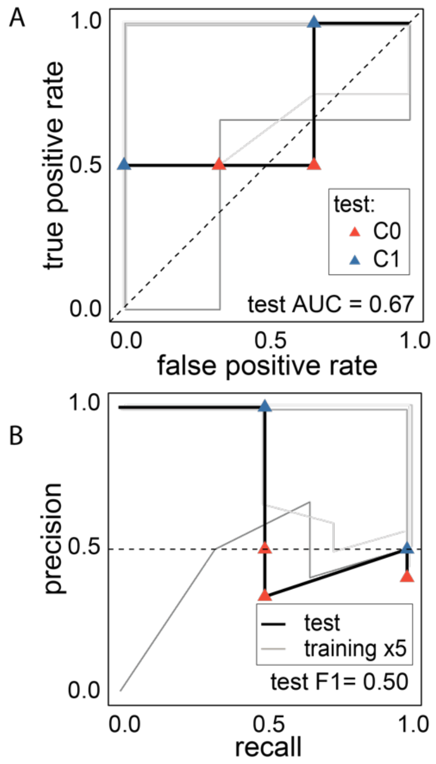

**Fig. S7.** BMM classifier performance on training and test data. **(A)** ROC and **(B)** PR curves of a random forest-based classifier trained on all eight conditions in the DiaMOND compendium. The model is tested with high-order combinations (4- and 5-drug combinations) that were excluded from training. Plots are labeled as in Fig. 3C. Training (black) and test (grey) performances are shown with lines. Test combinations are colored by outcome class as in (A). Performance metrics are shown on plots for training and test data (Area Under the ROC curve (AUC) and F1, harmonic mean of precision and recall). Dashed lines indicate theoretical "no-skill" model performance.

Fig. S8.

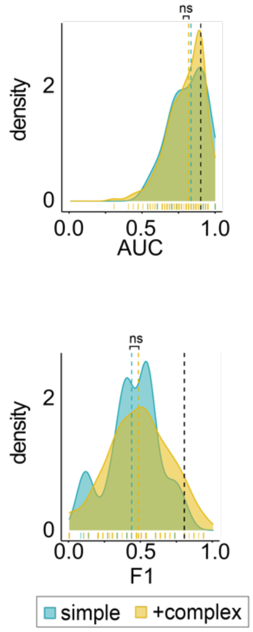

**Fig. S8.** BHeB *in vitro* model subset model performance distributions. Density distribution plots of estimated classifier performances from systematic survey of all possible *in vitro* model subsets. Distributions of **(A)** ROC AUC and **(B)** F1 are separated based on whether technically complex models (intracelluar, cholesterol-high, dormancy) are included (yellow) or whether only simple conditions (standard, acidic, butyrate, cholesterol, valerate) are considered. Colored dashed lines indicate the mean value for the distribution. The estimated performances when using all *in vitro* models is shown with black dashed lines. (Wilcoxon rank test: *** $p<0.005$. ** $p<0.01$. * $p<0.05$).

Fig. S9.

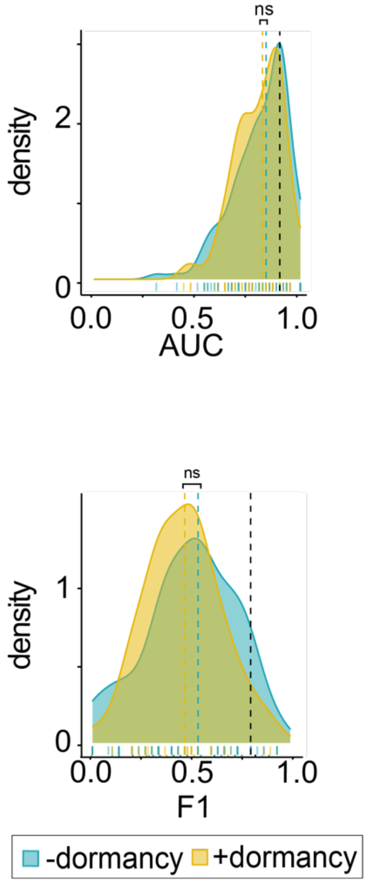

**Fig. S9.** BHeB *in vitro* model with and without dormancy model performance distributions. Density distribution plots of estimated classifier performances from a systematic survey of all possible *in vitro* model subsets. Distributions of **(A)** ROC AUC and **(B)** F1 are separated based on whether dormancy is included (yellow) or not. Colored dashed lines indicate the mean value for the distribution. The estimated performances when using all *in vitro* models are shown with black dashed lines. (Wilcoxon rank test: *** $p<0.005$. ** $p<0.01$. * $p<0.05$).

Table S1.

| Model | Estimated doubling time (days) | Measurement time (days) | Estimated relative doublings at each measurement |
|---|---|---|---|
| standard | 0.7 | 2.1, 2.7, 3.4, 4.2(CT) | 3, 4, 5, 6(CT) |
| intracellular | 1.5 | 2, 3, 4, 5(CT) | 1.3, 2, 2.8, 3.3(CT) |
| acidic | 2 | 6(C), 8, 10, 12(T) | 3(C), 4, 5, 6(T) |
| butyrate | 2 | 6(C), 8, 10(T) | 3(C), 4, 5(T) |
| valerate | 3 | 9(C), 12, 15(T) | 3(C), 4, 5(T) |
| cholesterol-high | 4 | 12(C), 16, 20, 24(T) | 3(C), 4, 5, 6(T) |
| cholesterol | 7 | 7(C), 14, 21, 28(T) | 1(C), 2, 3, 4(T) |
| dormancy | ND | 2(C), 3, 4(T) | 2.9(C), 4.3, 5.7(T) |

**Table S1.** Experiment time points and estimated growth amounts for *in vitro* models. *In vitro* models, estimated doubling times, experiment measurement times, and estimated relative doubling times for each model. Constant (C), terminal (T), and constant/terminal (CT) time points used in this study (as explained in the main text) are indicated. The dormancy model had no applicable growth rate (NA). Time points for dormancy were chosen based on the standard media growth rate added for the recovery period. Day 5 was chosen for the terminal time point for the intracellular model because the uninfected J774 cells began to lift from the plates.

Table S2.

| Drug name | 1 letter name | Cellular target process | Drug set |
|---|---|---|---|
| bedaquiline | B | ATP synthesis | DiaMOND compendium |
| clofazimine | C | respiratory inhibitor | DiaMOND compendium |
| ethambutol | E | cell wall synthesis | DiaMOND compendium |
| isoniazid | H | cell wall synthesis | DiaMOND compendium |
| linezolid | L | protein synthesis | DiaMOND compendium |
| moxifloxacin | M | DNA synthesis | DiaMOND compendium |
| pretomanid | Pa | cell wall synthesis/ nitric oxide production | DiaMOND compendium |
| pyrazinamide | Z | respiratory inhibitor | DiaMOND compendium |
| rifampicin | R | transcription | DiaMOND compendium |
| rifapentine | Rp | transcription | DiaMOND compendium |
| delamanid | D | cell wall synthesis/ nitric oxide production | Validation Drug |
| sutezolid | Su | protein synthesis | Validation Drug |
| SQ109 | Sq | cell wall synthesis | Validation Drug |
| gatifloxacin | G | DNA synthesis | Validation Drug |
| d-cycloserine | Cy | cell wall synthesis | Validation Drug |

**Table S2.** Drug information table. The drug set was either DiaMOND compendium (10 drugs used for systematic drug combinations) or test drugs (used for testing drug combinations predictions).

Table S3.

| Drug | butyrate | cholesterol | cholesterol-high | acidic | intracellular | dormancy | standard | valerate |
|---|---|---|---|---|---|---|---|---|
| bedaquiline | 0.07 | 0.07 | 0.06 | 0.26 | 0.08 | 0.87 | 0.77 | 0.01 |
| clofazimine | 1.89 | 2.15 | 1.53 | 1.64 | 2.30 | 3.99 | 2.90 | 0.28 |
| ethambutol | 7.93 | 7.89 | 6.52 | 49.19 | 56.65 | 14.62 | 2.24 | 16.99 |
| isoniazid | 0.02 | 0.04 | 0.02 | 0.05 | 0.13 | 0.26 | 0.08 | 0.03 |
| linezolid | 1.46 | 17.26 | 1.65 | 0.55 | 2.30 | 2.00 | 0.86 | 1.19 |
| moxifloxacin | 0.47 | 1.64 | 0.44 | 1.46 | 1.34 | 0.24 | 0.87 | 0.40 |
| pretomanid | 0.26 | 0.17 | 0.06 | 0.24 | 0.22 | 0.91 | 0.45 | 0.42 |
| pyrazinamide | 4394.83 | ND | 37.53 | 770.88 | 95.18 | ND | ND | 3461.45 |
| rifapentine | 0.20 | 0.09 | 0.26 | 0.04 | 0.53 | 0.06 | 0.02 | 0.12 |
| rifampicin | 0.169 | 0.276 | 0.368 | 0.191 | 1.213 | 0.060 | 0.087 | 0.183 |

**Table S3.** Drug $IC_{90}$ for *in vitro* models. Mean $IC_{90}$ (µg/mL) of single drugs calculated from fitted dose response curves (at least 2 replicates). $IC_{90}$ for individual replicates were calculated and then averaged. ND = not determined because $IC_{90}$ was not achieved.

Table S4.

| **Combination 1 letter** | **Drug number** | **Class** | **Combination class** | **References** |
|---|---|---|---|---|
| BL | 2 | C1 | Training | (*65*) |
| BPa | 2 | C1 | Training | (*65, 112*) |
| BZ | 2 | C1 | Training | (*113, 114*) |
| HE | 2 | C0 | Training | (*71*) |
| HZ | 2 | C0 | Training | (*71*) |
| HR | 2 | C0 | Training | (*71*) |
| RZ | 2 | C0 | Training | (*70, 71*) |
| BPaC | 3 | C1 | Training | (*114*) |
| BCZ | 3 | C1 | Training | (*23, 113, 114*) |
| BPaL | 3 | C1 | Training | (*64, 65*) |
| BZL | 3 | C1 | Training | (*64*) |
| BPaM | 3 | C1 | Training | (*112*) |
| BMZ | 3 | C1 | Training | (*23, 65, 115*) |
| BPaZ | 3 | C1 | Training | (*64, 112, 113*) |
| BRZ | 3 | C1 | Training | (*23*) |
| BZRp | 3 | C1 | Training | (*23, 114, 115*) |
| HRE | 3 | C0 | Training | (*116*) |
| HRM | 3 | C0 | Training | (*67*) |
| HMZ | 3 | C0 | Training | (*67*) |
| HRPa | 3 | C0 | Training | (*117*) |
| HRZ | 3 | C0 | Training | (*22, 23, 64, 66-71, 112, 114, 115, 117-121*) |
| HRpZ | 3 | C1 | Training | (*69, 118, 121*) |
| PaMZ | 3 | C1 | Training | (*22*) |
| MRPa | 3 | C0 | Training | (*22*) |
| MRZ | 3 | C0 | Training | (*22, 66-69*) |
| MRpZ | 3 | C1 | Training | (*69, 114, 121*) |
| RZPa | 3 | C0 | Training | (*22, 23, 117*) |

**Table S4.** Drug combinations with RMM outcomes for 2- and 3-way drug combinations. Single and 3 letter combination abbreviation along with classified score and with RMM outcome score, and literature reference.

Table S5.

| PC | U statistic | p |
|---|---|---|
| PC01 | 23 | 0.001 |
| PC27 | 142 | 0.007 |
| PC02 | 139 | 0.011 |
| PC04 | 131 | 0.034 |
| PC09 | 59 | 0.162 |
| PC03 | 60 | 0.178 |
| PC24 | 111 | 0.272 |
| PC23 | 66 | 0.294 |
| PC05 | 110 | 0.294 |
| PC08 | 66 | 0.294 |
| PC16 | 107 | 0.368 |
| PC26 | 107 | 0.368 |
| PC18 | 106 | 0.394 |
| PC20 | 105 | 0.422 |
| PC22 | 104 | 0.451 |
| PC17 | 102 | 0.512 |
| PC12 | 75 | 0.544 |
| PC06 | 76 | 0.577 |
| PC13 | 77 | 0.610 |
| PC15 | 77 | 0.610 |
| PC14 | 78 | 0.645 |
| PC11 | 81 | 0.753 |
| PC19 | 81 | 0.753 |
| PC25 | 95 | 0.753 |
| PC10 | 90 | 0.942 |
| PC21 | 86 | 0.942 |
| PC07 | 88 | 1.000 |

**Table S5.** RMM PC class separation. PCA transformed RMM classified 2- and 3-way data were used to compare C1 (16) and C0 (11) classified combinations. C1 and C0 combinations were compared for each PC separately using the Wilcoxon rank sum test. The U statistic and p-value are listed.

Table S6.

| Learning algorithm | AUC | F1 |
| --- | --- | --- |
| bart machine | 0.89 | 0.86 |
| random forest (RF) | 0.89 | 0.84 |
| xgboost | 0.73 | 0.75 |
| logistic regression | 0.68 | 0.78 |
| naïve bayes | 0.67 | 0.67 |
| support vector machine (SVM) | 0.65 | 0.48 |
| k-nearest neighbors (KNN) | 0.65 | 0.65 |

**Table S6.** Machine learning algorithm benchmarking performance metrics. PCA transformed RMM classified 2- and 3-way data were used to compare C1 and C0 classified combinations. Area Under the ROC curve (AUC) and F1 (harmonic mean of precision and recall).

Table S7.

| Combination 1 letter | Drug number | Class | Combination class | References |
|---|---|---|---|---|
| BCZE | 4 | C1 | 4-,5-way | (*120, 122*) |
| BPaLZ | 4 | C1 | 4-,5-way | (*64*) |
| BCZRp | 4 | C1 | 4-,5-way | (*114*) |
| BPaMZ | 4 | C1 | 4-,5-way | (*112*) |
| BMZRp | 4 | C1 | 4-,5-way | (*115*) |
| HRZC | 4 | C1 | 4-,5-way | (*123, 124*) |
| HRZE | 4 | C0 | 4-,5-way | (*71, 118-120, 123-126*) |
| HRpZE | 4 | C0 | 4-,5-way | (*118, 119, 127*) |
| MRZE | 4 | C1 | 4-,5-way | (*125, 127*) |
| HRZL | 4 | C0 | 4-,5-way | (*126*) |
| HRZM | 4 | C0 | 4-,5-way | (*67, 125, 127*) |
| HRZPa | 4 | C0 | 4-,5-way | (*117*) |
| RMZPa | 4 | C1 | 4-,5-way | (*22*) |
| HRZEC | 5 | C1 | 4-,5-way | (*119, 123*) |
| BSu | 2 | C1 | New Drug | (*113*) |
| BPaSu | 3 | C1 | New Drug | (*64, 113*) |
| BCZD | 4 | C1 | New Drug | (*128*) |
| BCZSq | 4 | C1 | New Drug | (*120, 128*) |
| HRZSu | 4 | C1 | New Drug | (*126*) |

**Table S7.** RMM model validation set. Drug combinations with RMM outcomes for selected higher order drug combinations and drug combinations that have new drugs. Single letter combination abbreviation along with classified score and with RMM outcome score, and literature reference.

Table S8.

| | Training performance | | Test performance | |
|---|---|---|---|---|
| ***in vitro* model subset** | **AUC** | **F1** | **AUC** | **F1** |
| all models | 0.92 | 0.84 | 0.75 | 0.86 |
| c | 0.91 | 0.90 | 0.75 | 0.61 |
| s | 0.91 | 0.82 | 0.69 | 0.70 |
| i | 0.90 | 0.79 | 1.00 | 0.40 |
| h | 0.83 | 0.67 | 0.85 | 0.94 |
| b | 0.79 | 0.73 | 0.72 | 0.74 |
| v | 0.79 | 0.68 | 0.80 | 0.77 |
| a | 0.68 | 0.81 | 0.58 | 0.91 |
| d | 0.67 | 0.76 | 0.50 | 0.80 |

**Table S8.** RMM single *in vitro* model classifier performance. Training performance for single *in vitro* model classifiers compared with the all-*in vitro* model classifier. PCA transformed RMM classified 2- and 3-way data were used to compare C1 and C0 classified combinations. Area Under the ROC curve (AUC) and F1(harmonic mean of precision and recall).

Table S9.

| **Combination 1 letter** | **Drug number** | **Class** | **Combination class** | **References** |
|---|---|---|---|---|
| B | 1 | C0 | Training | (*64, 65, 77, 129*) |
| L | 1 | C0 | Training | (*27, 28, 64*) |
| Pa | 1 | C0 | Training | (*64, 113, 117, 129*) |
| Rp | 1 | C0 | Training | (*28, 118*) |
| BL | 2 | C0 | Training | (*64, 65*) |
| BZM | 2 | C1 | Training | (*23, 65, 115*) |
| BPa | 2 | C0 | Training | (*64, 112, 129*) |
| BZ | 2 | C1 | Training | (*23, 77, 113*) |
| PaL | 2 | C0 | Training | (*64*) |
| PaM | 2 | C0 | Training | (*22*) |
| MZ | 2 | C0 | Training | (*22*) |
| PaZ | 2 | C1 | Training | (*22*) |
| RZ | 2 | C0 | Training | (*66, 70, 130*) |
| BPaC | 3 | C1 | Training | (*114*) |
| BZC | 3 | C1 | Training | (*23*) |
| BPaL | 3 | C1 | Training | (*64, 65*) |
| BZL | 3 | C1 | Training | (*23*) |
| BPaM | 3 | C1 | Training | (*65, 112*) |
| BPaZ | 3 | C1 | Training | (*23, 64, 112*) |
| BRZ | 3 | C1 | Training | (*23*) |
| BZRp | 3 | C1 | Training | (*23, 115*) |
| HRE | 3 | C0 | Training | (*131*) |
| HZM | 3 | C0 | Training | (*67*) |
| HRM | 3 | C0 | Training | (*67*) |
| HZPa | 3 | C0 | Training | (*117*) |
| HRPa | 3 | C0 | Training | (*117*) |
| HRZ | 3 | C0 | Training | (*22, 64, 66-70, 112-115, 117-* |

| | | | | |
|---|---|---|---|---|
| | | | | *120, 129, 130*) |
| HRpZ | 3 | C1 | Training | (*118, 119, 121*) |
| PaMZ | 3 | C1 | Training | (*112*) |
| MRPa | 3 | C0 | Training | (*22*) |
| MRZ | 3 | C1 | Training | (*22, 66-69, 121, 130*) |
| MRpZ | 3 | C1 | Training | (*69, 121*) |
| RZPa | 3 | C1 | Training | (*22, 23, 117*) |
| BPaLZ | 4 | C1 | 4-,5-way | (*64*) |
| BPaMZ | 4 | C1 | 4-,5-way | (*65, 112*) |
| BZRpM | 4 | C1 | 4-,5-way | (*115*) |
| HRZC | 4 | C0 | 4-,5-way | (*123, 124*) |
| HRZE | 4 | C0 | 4-,5-way | (*118-120, 123-125*) |
| HRpZE | 4 | C1 | 4-,5-way | (*118, 119*) |
| MRZE | 4 | C1 | 4-,5-way | (*125*) |
| HRZM | 4 | C0 | 4-,5-way | (*67, 125*) |
| HRZPa | 4 | C0 | 4-,5-way | (*117*) |
| MRZPa | 4 | C1 | 4-,5-way | (*22*) |
| HRZEC | 5 | C0 | 4-,5-way | (*119, 123*) |
| BCZE | 4 | C1 | 4-,5-way | (*120, 122*) |
| HRZL | 4 | C0 | 4-,5-way | (*126*) |
| BCZD | 4 | C1 | New Drug | (*128*) |
| BCZSq | 4 | C1 | New Drug | (*120, 128*) |
| BPaSu | 3 | C1 | New Drug | (*113*) |
| BSu | 2 | C1 | New Drug | (*113*) |
| CyM | 2 | C0 | New Drug | (*132*) |
| D | 1 | C0 | New Drug | (*113*) |
| GR | 2 | C0 | New Drug | (*133*) |
| HRZSq | 4 | C1 | New Drug | (*134*) |
| HRZSu | 4 | C1 | New Drug | (*126*) |
| HRSq | 3 | C0 | New Drug | (*134*) |
| PaSu | 2 | C1 | New Drug | (*64*) |
| Su | 1 | C0 | New Drug | (*64*) |

**Table S9.** Drug combinations with BMM outcomes for 1-, 2- and 3-way drug combinations. Single letter combination abbreviation along with classified score and with BMM outcome score, and literature reference.

Table S10.

| **Combination 1 letter** | **Drug number** | **Class** | **Combination class** | **References** |
|---|---|---|---|---|
| H | 1 | C0 | Training | (*27*) |
| L | 1 | C0 | Training | (*27, 28*) |
| Z | 1 | C0 | Training | (*27, 28, 77, 129*) |
| R | 1 | C0 | Training | *26, 27, 69, 110*) |
| Rp | 1 | C0 | Training | (*28, 118*) |
| BZ | 2 | C0 | Training | (*77*) |
| MZ | 2 | C0 | Training | (*75*) |
| RZ | 2 | C0 | Training | (*129*) |
| BMZ | 3 | C1 | Training | (*65*) |
| HRE | 3 | C0 | Training | (*131*) |
| HRZ | 3 | C0 | Training | (*75*) |
| MRpZ | 3 | C1 | Training | (*75*) |
| BPaMZ | 4 | C1 | Training | (*65*) |
| HRZE | 4 | C0 | Training | (*119, 120, 125, 131*) |
| MRZE | 4 | C1 | Training | (*125*) |
| HRZM | 4 | C1 | Training | (*125*) |

**Table S10.** Drug combinations with BHeB outcomes for 1-, 2- and 3-way drug combinations. Single letter combination abbreviation along with classified score and with BHeB outcome score, and literature reference.

Table S11.

| PC | U statistic | p |
|---|---|---|
| PC03 | 55 | 0.001 |
| PC11 | 41 | 0.145 |
| PC14 | 41 | 0.145 |
| PC07 | 37 | 0.320 |
| PC13 | 19 | 0.377 |
| PC08 | 19 | 0.377 |
| PC15 | 21 | 0.510 |
| PC16 | 22 | 0.583 |
| PC05 | 32 | 0.661 |
| PC09 | 32 | 0.661 |
| PC10 | 24 | 0.743 |
| PC01 | 25 | 0.827 |
| PC02 | 25 | 0.827 |
| PC12 | 26 | 0.913 |
| PC06 | 29 | 0.913 |
| PC04 | 28 | 1.000 |

**Table S11.** BHeB PC class separation. PCA transformed BHeB classified 1-, 2- and 3-way data to compare C1(5) and C0(11) classified combinations. C1 and C0 combinations were compared for each PC separately using the Wilcoxon rank-sum test. The U statistic and p-value are listed.
